# Enhancing and inhibitory motifs have coevolved to regulate CD4 activity

**DOI:** 10.1101/2021.04.29.441928

**Authors:** Mark S. Lee, Peter J. Tuohy, Caleb Kim, Katrina Lichauco, Heather L. Parrish, Koenraad Van Doorslaer, Michael S. Kuhns

## Abstract

CD4^+^ T cells use T cell receptor (TCR)-CD3 complexes, and CD4, to respond to peptide antigens within MHCII molecules (pMHCII). We report here that, through ∼435 million years of evolution in jawed vertebrates, purifying selection has shaped motifs in the extracellular, transmembrane, and intracellular domains of eutherian CD4 that both enhance pMHCII responses and are coevolving with residues in an intracellular motif that inhibits pMHCII responses. Importantly, while CD4 interactions with the Src kinase, Lck, are classically viewed as the key determinant of CD4’s contribution to pMHCII responses, we found that without the inhibitory motif CD4-Lck interactions are not necessary for robust responses to pMHCII. In summary, motifs that mediate events on the outside and inside of CD4^+^ T cells coevolved to finetune the relay of pMHCII-specific information across the membrane. These results have implications for the evolution and function of complex transmembrane receptors and for biomimetic engineering.

## INTRODUCTION

The immunological ‘Big Bang’ that gave rise to RAG-based antigen receptor gene rearrangement in jawed vertebrates produced an adaptive immune system in which each naïve B and T cell expresses a clonotypic B or T cell receptor (BCR or TCR) encoded by gene segments rearranged by Rag-1 and Rag-2 (Bernstein et al., 1996). Most T cells use their TCRs together with either CD4 or CD8 coreceptors to recognize peptide antigens presented by MHCII or MHCI (pMHCII or pMHCI) molecules. Interestingly, *Callorhinchus milii*, a cartilaginous fish with an immune system that is thought to resemble the primordial adaptive immune system, encodes BCRs, TCRs, MHCI, MHCII, and CD8 within its genome – but an orthologous gene encoding CD4 appears to be absent, as are genes for proteins associated with CD4^+^ T cell helper (Th) or regulatory (Treg) functions (e.g. *FoxP3* and *Rorc*) (Venkatesh et al., 2014). These findings were taken as evidence that CD4 evolved to help direct more elaborate T cell functions than could be achieved by TCR engagement of pMHCII alone.

While this idea is intriguing, most models attribute the quantity and quality of pMHCII-specific signaling to the biophysical properties of TCR-pMHCII interactions; they then assign CD4 a supporting role whereby it interacts with the Src kinase, Lck, via an intracellular CQC clasp motif to recruit Lck to phosphorylate the immunoreceptor tyrosine-based activation motifs (ITAMs) of the TCR-associated CD3 signaling modules when both the TCR and CD4 coincidently engage pMHCII (Glaichenhaus et al., 1991; Guy and Vignali, 2009; Kuhns and Davis, 2012; Malissen and Bongrand, 2015; Turner et al., 1990; Yin et al., 2012). It is challenging to envision how CD4 might direct elaborate pMHCII responses based on these models.

However, more recent data suggest that CD4 and the TCR-CD3 complex work in concert to mediate pMHCII responses. CD4’s extracellular domain (ECD) can increase TCR dwell time on pMHCII and position its intracellular domain (ICD) in a defined relationship with the TCR-CD3 complex (Glassman et al., 2016; Glassman et al., 2018; Guy and Vignali, 2009). Furthermore, CD4 molecules that are not associated with Lck have been proposed to compete with those that are to limit the number of TCR-CD3 complexes phosphorylated by Lck, thus setting a threshold for the duration of TCR-pMHCII interactions required to initiate signaling (Stepanek et al., 2014). These models help explain how the stability and composition of TCR-CD3-pMHCII-CD4(+/-Lck) assemblies can influence the quantity of ITAM phosphorylation. But, due to their focus, they did not consider evidence that motifs in the transmembrane domain (TMD) and ICD aside from the CQC clasp may also regulate CD4 function (Fragoso et al., 2003; Parrish et al., 2015; Popik and Alce, 2004).

As nature has been experimenting with CD4 in a variety of jawed vertebrate species for ∼435 million years, we performed evolution reconstruction analyses of CD4 homologs to gain insights into the results of those experiments and better understand CD4 function. Our analyses of extant CD4 orthologs from fish, reptiles, birds, and mammals identified five putative motifs within the TMD and ICD that are unique to mammalian CD4, or found only in eutherians (placental mammals), and contain residues under purifying selection. Follow-on biochemical and functional studies revealed that most of these motifs enhance mouse CD4 function by influencing the membrane domains in which CD4-Lck interactions occur, while one of these motifs inhibits the function of CD4 molecules that are not associated with Lck. Further analyses revealed a network of residues in motifs in the ECD, TMD, and ICD that enhance CD4 activity that are coevolving with residues in the inhibitory motif in the ICD. These data provide evidence that CD4 is under evolutionary pressure to finetune the relay of pMHCII-specific information across the membrane by coordinating events that happen on the outside and inside of the cell. The findings have broad implications for how multi-subunit transmembrane receptors relay ligand-specific information across the cell membrane, as well as for biomimetic engineering of synthetic receptors.

## RESULTS

### Evolutionary analysis of CD4

We performed multiple analyses of available vertebrate CD4 ortholog sequences, representing ∼435 million years of evolution, to understand how ancient and ongoing environmental challenges have influenced CD4. We used mouse CD4 (numbering by UniProt convention) as a reference to facilitate comparisons between evolutionary analyses and experimental studies. An analysis of sequence conservation between the full set of extant CD4 molecules, or only mammalian CD4 molecules, showed particular conservation in the ICD that suggested more than just the CQC clasp is important for CD4 function (**Figure S1A and S1B**)(Capra and Singh, 2007). We also performed fixed effects likelihood (FEL) analysis to determine nonsynonymous (dN) and synonymous (dS) substitution rates within the CD4 coding sequence to formally test the importance of each residue in both the full and mammalian only data sets (Kosakovsky Pond and Frost, 2005). By this analysis, positions under diversifying selection have a dN:dS ratio >1 and those under purifying selection have a dN:dS ratio <1 (**Figure S1C and S1D**). Of the 17 codons under diversifying selection, 16 (94.1%) are distributed across the CD4 ECD while only one is found in the TMD. Of the 126 residues under purifying selection, 98 are distributed across the CD4 ECD (24.8% of all codons in the ECD). In contrast, 45.5% of TMD codons (10 of 22) and 45% (18 of 40) within the ICD are under purifying selection. In sum, the ECD contains residues under selective pressure due either to their structural or functional importance, while the TMD and ICD of mammalian CD4 are particularly constrained due to their importance. As the ICD of human CD4 is largely unstructured, the residues in the ICD are likely to be functionally important (Kim et al., 2003).

We proceeded to focus on an ECD region that includes the D3 domain solvent-exposed nonpolar patch that enhances CD4 function, the TMD, and the ICD to more closely interrogate their evolution (Glassman et al., 2018). Accordingly, we generated a maximum likelihood phylogenetic tree that included the predicted most recent common ancestor (MRCA) sequences at each node to evaluate the evolutionary events that dictate structure and function at our regions of interest (**Figure 1A** **and Figure S1E**)(Hochberg and Thornton, 2017). We also evaluated residue occurrence in eutherians by logo plot analysis to better consider positional variability of residues within this clade. Additionally, we related evolutionary changes on a residue-by-residue basis to our FEL results for all extant CD4 molecules, as well as the mammalian only dataset, to gain a more complete picture of the pressures shaping CD4 molecules (**Figure 1B and 1C**). Particular attention was given to mouse and human CD4 due to their experimental and human health relevance.

**Figure 1.**
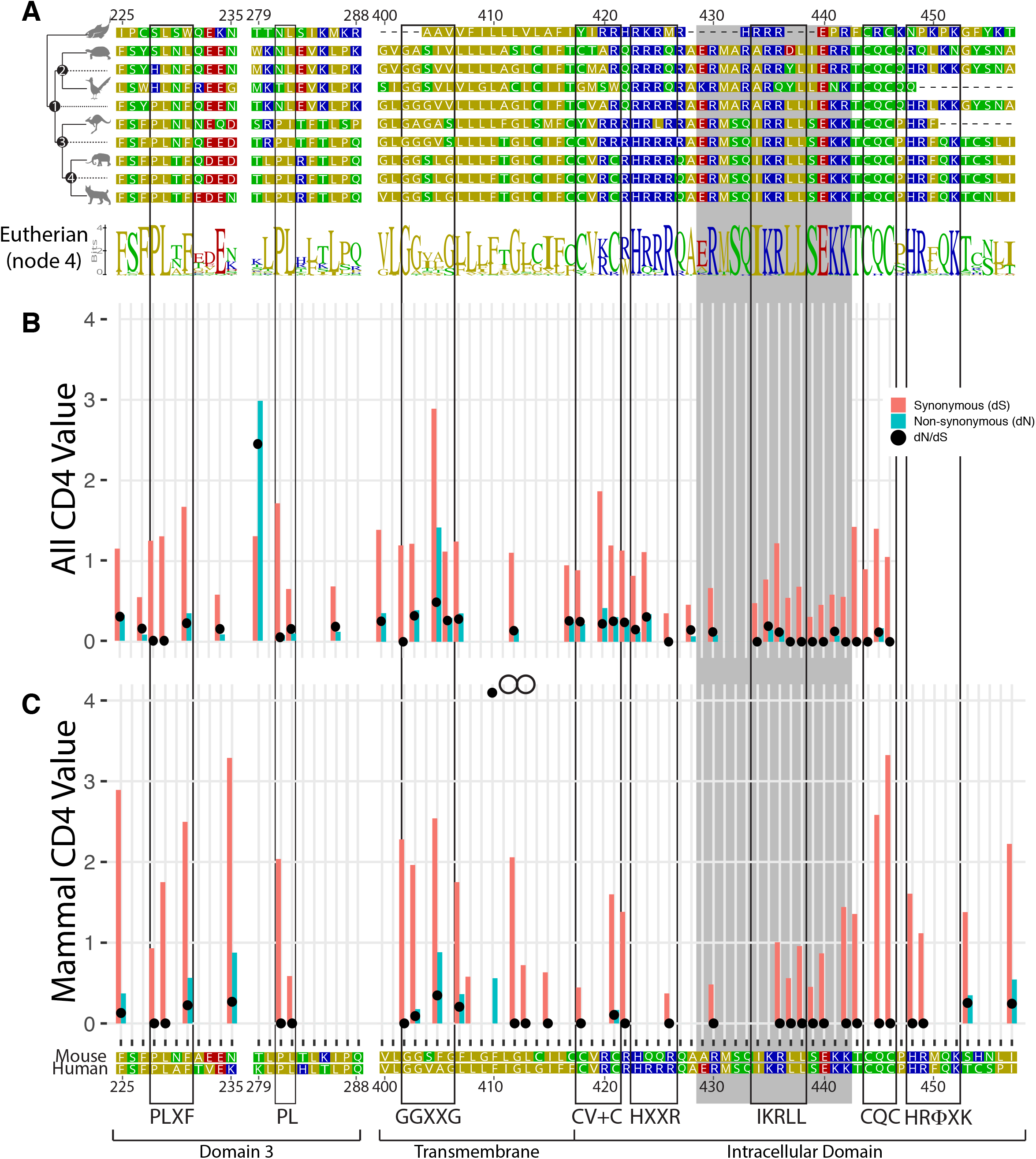
Evolutionary analysis of the CD4 molecule. (A) Reduced representation maximum likelihood phylogenetic tree clusters of CD4 sequences are shown with mouse CD4 numbering (uniprot) used as a reference. Residues are colored based on sidechain polarity with dashes (-) indicating an evolutionary insertion or deletion event. Predicted most recent common ancestor (MRCA) sequences are shown at each node in the tree (Node 1-4). Logo plots of extant eutherian CD4 sequences are aligned at the bottom of the tree in which each stack of letters represents the sequence conservation at that position in the alignment and the height of symbols indicates the relative frequency of each amino acid at that specific position. (B) Synonymous (dS, red bars) and nonsynonymous (dN, blue bars) substitution rates within the CD4 coding sequence are shown as calculated for all CD4 orthologs included in the initial phylogenetic analysis using the Fixed Effects Likelihood (FEL) method. Only bars for which the likelihood ratio test indicated statistical significance (alpha = 0.1) are shown. Black circles show the ratio of both these values. Codons under diversifying selection have a dN:dS ratio >1 and those under purifying selection have a dN:dS ratio <1. (C) dS and dN substitution rates are shown as in (B) for the dataset containing only mammalian CD4 sequences to include the region C-terminal of the CQC clasp that is highly variable between phylogenetic clades, and could not be confidently aligned, but is highly conserved in mammals. The codon corresponding to F410 had a dS of 0, leading to a dN/dS ratio of infinity. Human and mouse CD4 sequences are aligned at the bottom for reference. Boxes are used to highlight motifs discussed in this study, while the grey shading indicates the helix-turn region within the ICD. See also Figure S1.

First, we focused on the evolutionary history of the PLXF motif (mouse 228-231) and P281 residue in the D3 domain of the CD4 ECD to ask if our analyses would highlight residues that are adjacent in 3D space, form a solvent-exposed nonpolar patch, facilitate the formation and stability of TCR-CD3-pMHCII-CD4 assemblies, and increase responses to weak and agonist pMHCII (Glassman et al., 2018; Wu et al., 1997). The MRCA of all amniotes contains a PLXF motif (mouse 228-231) in the D3 domain and is fixed in mammals (**Figure 1A**, node 1). P281 is not found in the amniote MRCA but is fixed in all mammals, suggesting that it arose later. Importantly, P228, L229, F231, and P281 all have very small dN:dS values that are primarily driven by low dN rates, suggesting that they are of structural or functional importance. Indeed, structural analysis indicates that L229 is buried in the hydrophobic core of the D3 domain as is L282 adjacent to P281, implying a structural function, while the solvent exposed P228, F231, and P281 are known to be important for CD4 function, providing a possible explanation for why these residues are under purifying selection (Glassman et al., 2018; Wu et al., 1997).

We next turned our attention to residues within the TMD and ICD by focusing on motifs that: 1) displayed a stepwise refinement from ancestral sequences towards extant sequences; 2) contain highly conserved residues with evidence of purifying selection. For example, the MRCA of all amniotes (node 1) contains a GG patch (G402 and G403) in its TMD that is lost in non-avian reptiles but present in extant birds (**Figure 1A**). In the eutherian MRCA (node 4) the GG patch expanded into a highly conserved GGXXG motif that is present in the majority of eutherian CD4 proteins (see logo plot) including mouse and human. The selection analysis also identified G402 and G403 using both the full and mammal-only dataset, while G406 was only identified in the full dataset (**Figure 1B and 1C**). As we have ruled out a role for the GGXXG motif in mediating mouse CD4 dimerization, it is likely to mediate heterotypic protein-protein or protein-cholesterol interactions (Parrish et al., 2015; Teese and Langosch, 2015).

Using our approach we also identified a CV+C motif (“+” represents a basic residue) that fixated in eutherians (**Figure 1A-1C**). Palmitoylation of these cysteines is reported to influence membrane raft localization, although this is controversial as association with Lck may also localize CD4 to membrane rafts (Crise and Rose, 1992; Fragoso et al., 2003; Ladygina et al., 2011). Our analysis suggests that the combination of the CV+C and the GGXXG/S motifs are unique to extant eutherians, co-arose during evolution, and may work together to regulate CD4 membrane localization.

A poly-basic RHRRR motif in the juxtamembrane region of the ICD was also previously reported to impact human CD4 localization to membrane rafts (Popik and Alce, 2004). Our comparative analyses suggest a core HXXR motif (mouse 423-426) with H423 and R426 under purifying selection (**Figure 1A-1C**).

Further downstream, NMR has shown that the ICD of human CD4 contains a helix-turn structure, the sequence of which is highly conserved in all mammalian CD4 molecules (grey shaded region, 429-442, **Figure 1A**)(Kim et al., 2003; Willbold and Rosch, 1996). A conserved IKRLL motif, resembling known I/LXXL protein-interaction motifs, is embedded within the helix. Its origins are evident in the amniote MRCA (node 1) via the presence of a dileucine repeat. Reptiles and birds diverged away from this motif, while the MRCA of mammals (node 3) evolved a conserved IXRLL. This region includes a number of residues under purifying selection within the full dataset analysis (R430, 434-438 that make up the IKRLL motif, S439, and 440-442 that form the turn) or within the more limited mammalian dataset (436-440 and 442) (**Figure 1B and 1C)**. Many of these residues are important for the helix-turn structure, while I434, L437, and L438 are reported to regulate CD4-Lck interactions and endocytosis, suggesting that these residues are under purifying selection due to their role in a multi-functional hub (Kim et al., 2003; Sleckman et al., 1992). Intriguingly, the HIV-1 Nef protein interacts with the IKRLL motif to promote CD4 downregulation (Kwon et al., 2020). Such host-pathogen interactions often leave evidence of diversifying selection in host genes yet, in this case, it is possible that HIV-1 Nef targets the IKRLL motif due to its functional necessity, limiting the ability of CD4 to ‘evolve around’ Nef binding (Daugherty and Malik, 2012; Elde et al., 2009).

Finally, C-terminal to the CQC clasp, mammalian CD4 contains a consensus HRΦQK motif (mouse 448-452 in which Φ represents a large hydrophobic residue; **Figure 1A**). This putative motif is not present in extant fish, reptiles, birds, or even the marsupial CD4 orthologues sequenced to date. Yet, within the mammalian dataset, the codons for H448 and R449 were found to be under purifying selection (**Figure 1C**). The NMR solution structure of the CD4 ICD indicates that this region is unstructured within human CD4, thus these residues are likely to be of functional importance (Kim et al., 2003).

To summarize, our analyses highlight motifs in the ECD, TMD, and ICD of extant eutherian CD4 that contain residues under purifying selection, indicating that they are important for CD4 structure and/or function. We therefore proceeded to investigate how residues within these motifs impact CD4 membrane domain localization, CD4-Lck association, and pMHCII-responses.

### The GGXXG and CV+C motifs together impact membrane domain localization and function

We first tested the hypothesis that together the GGXXG and CV+C influence membrane domain localization and function using previously established approaches (Fragoso et al., 2003; Parrish et al., 2016). We included the CQC clasp because Lck has myristylation and palmitoylation sites that could influence membrane domain localization of CD4 when associated via the clasp (Ladygina et al., 2011). 58α^-^β^-^ cells, which lack expression of endogenous TCR and CD4, were generated expressing the 5c.c7 TCR, which recognizes the 88-103 moth cytochrome c peptide (MCC) presented by mouse MHCII I-E^k^, along with CD4 (WT), a GGXXG to GVXXL mutant (TMD), a CV+C to SV+S mutant (Palm), a CQC to SQS clasp mutant (Clasp), a double TMD+Palm (TP) mutant, or a triple TMD+Palm+Clasp (TPC) mutant for functional analysis (**Figure S2A**)(Letourneur and Malissen, 1989).

To measure membrane domain localization and Lck association we performed sucrose gradient fractionation of detergent lysates. Proteins that float on the gradient localize to detergent resistant membrane domains (DRMs) rich in membrane raft components such as cholesterol and sphingolipids (Pike, 2006). The remaining proteins localize to detergent soluble membrane domains (DSMs). CD4 molecules in each fraction were immunoprecipitated (IP’d) onto polystyrene beads and stained with anti-CD4, the cholera toxin B-subunit (CTxB) that binds GM1 gangliosides associated with particular membrane rafts, and anti-Lck. Quantification of signal intensity was performed by flow cytometry (**Figure S2B-S2D**).

We focused on the percent of CD4 signal in each fraction, relative to the total, to account for any differences in the amount of CD4 between samples and independent experiments (**Figure 2A**). Area under the curve (AUC) analysis of the DRMs and DSMs revealed that Palm and Clasp mutant localization to the DRMs was lower than WT in our sample size, while DRM localization for the TP and TPC mutants was further reduced. These data indicate that the CQC and CV+C motifs, and particularly the GGXXG plus CV+C together, mediate CD4 localization to the DRMs. The TP and TPC showed slight increases in localization to the DSMs.

**Figure 2.**
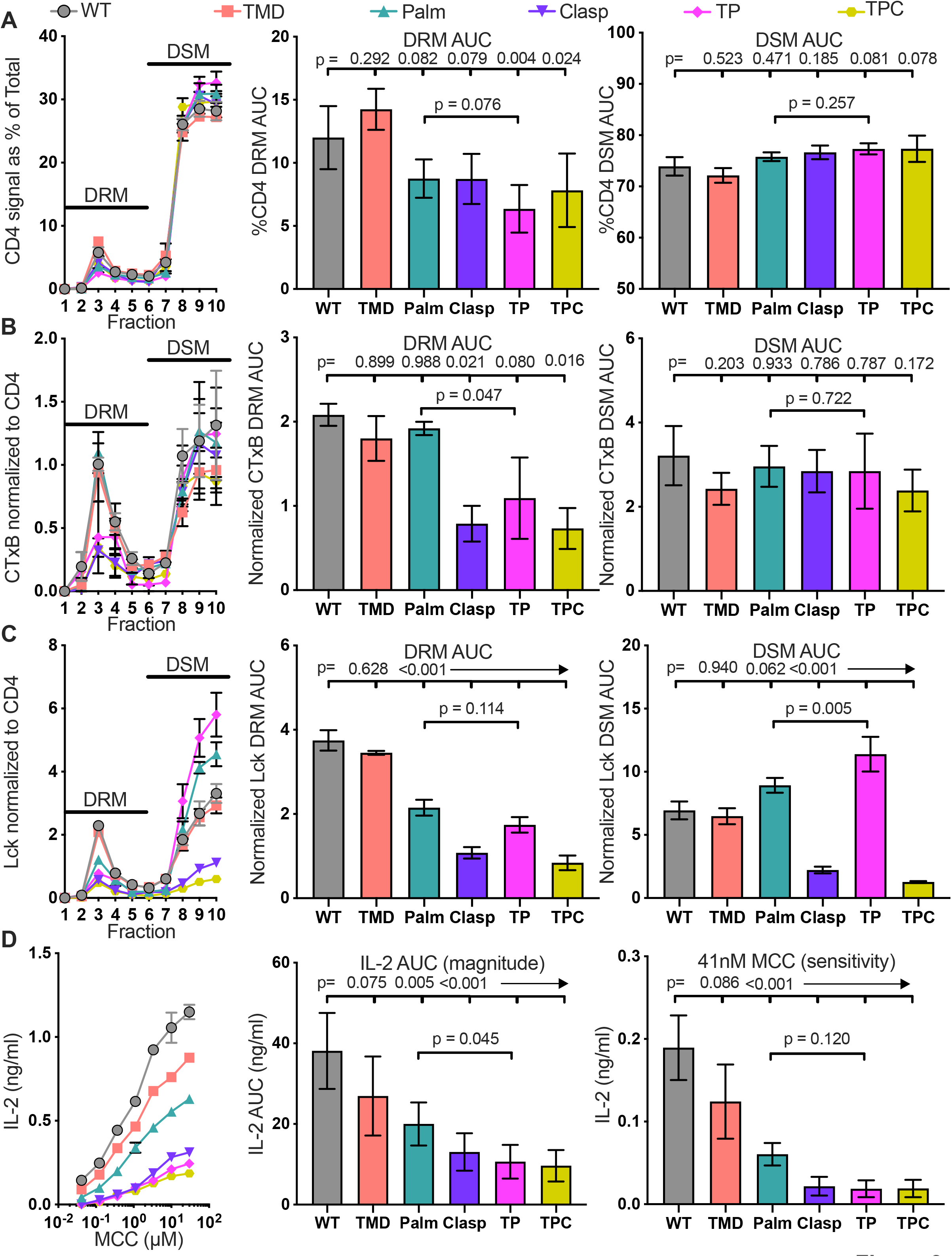
The GGXXG + CV+C motifs influence CD4 membrane domain localization and function. (A) CD4 signal for each sucrose gradient fraction is shown as a percent of the total CD4 signal detected in all fractions (left). The area under the curve (AUC) is presented for the normalized CD4 signal in the detergent resistant membrane (DRM) fractions (center) and detergent soluble membrane (DSM) fractions (right). (B) Cholera toxin B (CTxB) signal is shown for each sucrose fraction normalized to the CD4 signal detected in the corresponding fraction (left). The AUC is shown for the normalized CTxB signal in the DRM (center) and DSM fractions (right). (C) Lck signal is shown for each sucrose fraction normalized to the CD4 signal detected in the corresponding fraction (left). The AUC is shown for the normalized Lck signal in the DRM (center) and DSM fractions (right). (D) IL-2 production is shown in response to a titration of MCC peptide (left). AUC analysis for the dose response is shown as a measure of the response magnitude (center). The average response to a low dose (41nM) of peptide is shown as a measure of sensitivity (right). For (A-C) each data point represents the mean +/- SEM for the same three independent experiments (biological replicates). For (D), the dose response represents one of three experiments showing the mean +/- SEM of triplicate wells (technical replicates). The magnitude and sensitivity data represent the mean +/- SEM of three independent experiments (biological replicates). One-way ANOVA with a Dunnett’s posttest for comparisons with WT samples, or a Sidak’s posttest for comparisons between selected samples, were performed. See also Figure S2.

We next normalized the CTxB signal in each fraction to the CD4 signal in that fraction to assess the amount of GM1 that co-IP’d with CD4 per fraction (**Figure 2B**). AUC analysis revealed that the Clasp, TP, and TPC mutants within the DRMs had reduced staining with CTxB, and the TP mutant had lower CTxB staining than the Palm mutant. These data indicate that, for CD4 molecules within the DRM, the clasp influences CD4 association with GM1-containing rafts and the GGXXG and CV+C residues together do the same.

We also normalized the Lck signal in each fraction to the CD4 signal detected in that fraction to analyze the amount of Lck that co-IP’d with CD4 per fraction (**Figure 2C**). The Palm, Clasp, TP and TPC signals were all greatly reduced for AUC analysis of the DRM, indicating that both the CV+C and clasp motifs influence association with Lck in the DRM. AUC analysis also showed that only the Clasp and TPC mutants had reduced Lck association in the DSM, whereas the Palm and particularly the TP mutant had increased association with Lck relative to the WT or T mutant alone. These data indicate that the clasp controls if CD4 interacts with Lck, while the GGXXG and CV+C motifs together influence the type of membrane domain in which Lck-associated CD4 molecules localize.

IL-2 production was measured in response to a titration of MCC peptide to determine how these motifs influence pMHCII responses. If the frequency of CD4-Lck interactions is the chief determinant for pMHCII responses then only the Clasp and TPC mutants should reduce IL-2 production as the Palm and TP mutants interacted with Lck in the DSM (Glaichenhaus et al., 1991; Stepanek et al., 2014). But, if CD4 association with Lck in the DRMs is important, then the Palm and TP mutants would be expected to have reduced IL-2 production. We observed a hierarchy of IL-2 production of WT > TMD > Palm > Clasp ≥ TP ≥ TPC in response to a titration of MCC (**Figure 2D**), with AUC analysis showing that the magnitude of the response across the peptide titration was lower for each mutant cell line compared with the WT. The TP mutant also produced less IL-2 than the Palm mutant. Finally, we observed the same hierarchy of IL-2 production in response to the lowest dose of MCC tested (41nM). These data suggest the CV+C and GGXXG motifs that co-arose in evolution work together to enhance pMHCII responses by localizing Lck-associated molecules in the DRM.

### The HXXR and HRΦQK motifs influence membrane domain localization and association with Lck

We next analyzed the HXXR and HRΦQK motifs which both contain core basic residues under purifying selection. 58α^-^β^-^ cells expressing a R426A mutant of the HXXR motif, a R449A or a K452A mutant of the HRΦQK motif, or a R449A.K452A double mutant (Dbl) were generated to explore how these residues work individually or in combination. Each mutation reduced CD4 surface expression and CD4 signal in the sucrose gradients relative to WT, suggesting that these residues influence CD4 expression or turnover (**Figure S3A**). While the CTxB staining was unremarkable, the raw Lck signal was interesting when considered with CD4 levels because, while R426A and R449A had lower Lck signal in the DRMs, the signal was elevated in the DSMs compared with WT (**Figure S3B and S3C**). For these mutants we therefore calculated the total AUC for the entire sucrose fractionation curve for raw Lck signal (**Figure S3D**), which yielded a measure of the total Lck signal co-IP’d with the mutant CD4s that was not different from the WT (Mean ± SEM: WT = 2628 ± 208; R429A = 2921 ± 377; R449A = 2645 ± 430; K452A = 2385 ± 267; Dbl = 2155 ± 198; ns by One-way ANOVA with Dunnett’s posttest for mutants compared with WT), suggesting a higher overall frequency of CD4-Lck association in the mutant cells.

Analysis of normalized sucrose gradient data revealed that R426A and R449A reduced CD4 localization to the DRM, while K452A and the Dbl mutant had less of an impact. The R426A mutant also increased localization to the DSM (**Figure 3A**). The CTxB signal associated with CD4 in the DRMs or DSMs for these mutants was variable (**Figure 3B**). Similarly, the Lck signal normalized to the CD4 signal for the DRMs was unremarkable, but the R426A and R449A mutants were associated with more Lck signal in the DSMs than the WT (**Figure 3C**). The K452 mutant had lower levels of associated Lck than the R449A mutant in the DSM and the double mutant mirrored the phenotype of the K452A mutant, indicating that the R449 and K452 residues are not equivalent in the HRΦQK motifs. Together, the data indicate that the HxxR and HRΦQK motifs influence CD4 membrane domain localization and association with Lck therein.

**Figure 3.**
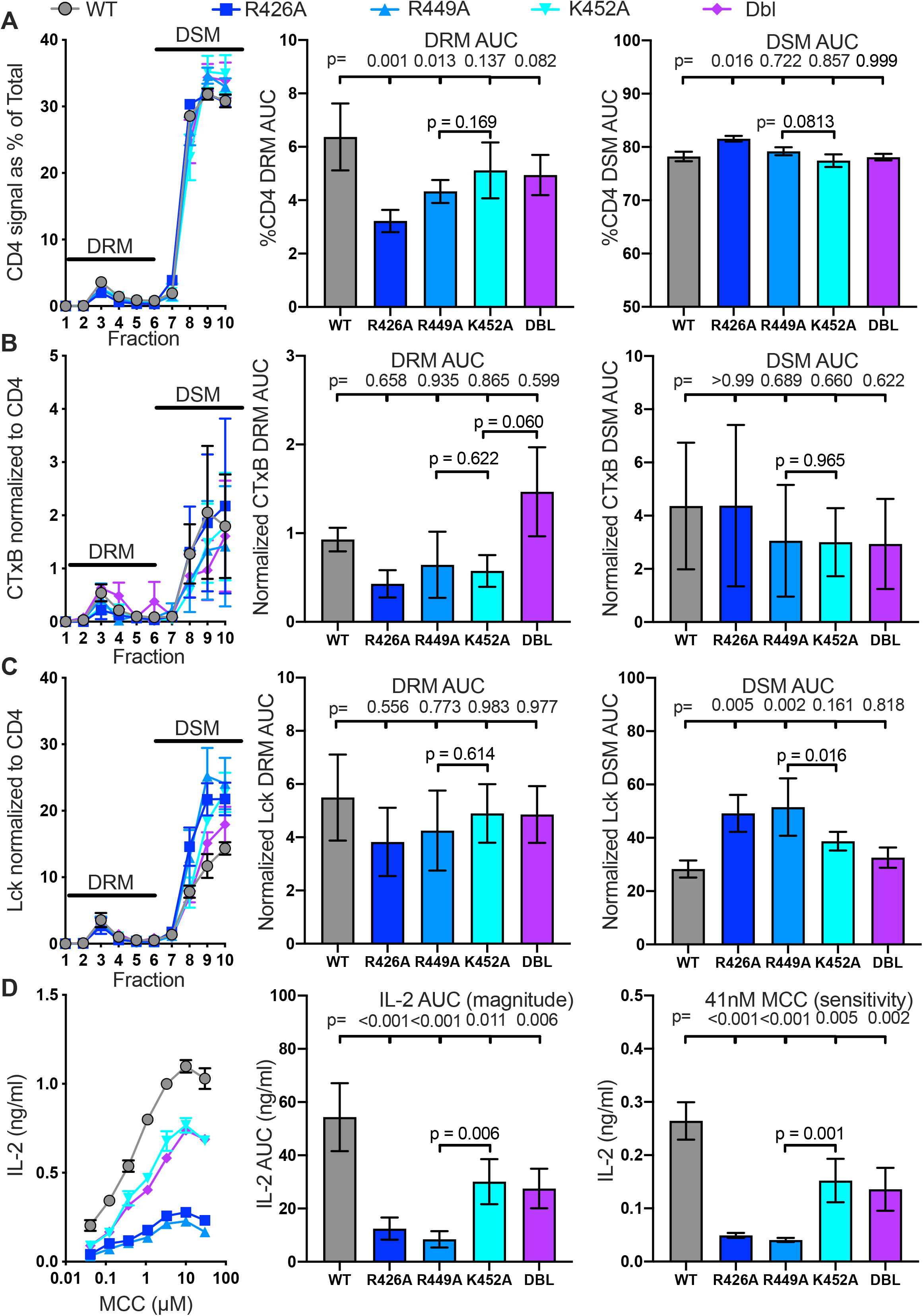
The HXXR and HRΦQK motifs influence CD4 membrane domain localization and function. (A) CD4 signal normalized as a percent of the total is shown for each sucrose gradient fraction (left) along with AUC analysis for the DRM (center) and DSM fractions (right). (B) Cholera toxin B (CTxB) signal normalized to CD4 signal detected is shown for each sucrose fraction (left) along with the AUC analysis for the DRM (center) and DSM fractions (right). (C) Lck signal normalized to CD4 signal is shown for each sucrose fraction (left) along with the AUC analysis for the DRM (center) and DSM fractions (right). (D) IL-2 dose response to MCC peptide (left), AUC analysis as a measure of the response magnitude (center), and the average response to a low dose (41nM) of MCC as a measure of sensitivity (right) are shown. For (A-D) the data are presented as in Figure 2. One-way ANOVA was performed with a Dunnet’s posttest for comparisons with WT samples and a Sidak’s posttest for comparisons between selected samples. See also Figure S3.

Here again, if the frequency of CD4-Lck association alone determines pMHCII-responses then the mutants should have equivalent pMHCII-responses to the WT. But AUC analysis showed that each mutant cell line produced less IL-2 than the WT (**Figure 3D**). These data are consistent with the idea that the membrane domain in which CD4 interacts with Lck is important for pMHCII responses.

### The intracellular helix interacts with Lck and attenuates pMHCII responses

Focusing on the intracellular helix-turn structure of the ICD, we replaced residues 430-442 with NGPGGNPGGNAGG to disrupt the chemical and structural nature of the helix-turn region but maintain its length. We also combined this helix (H) mutant with the clasp mutant (HC) to explore how they work together. The H and HC mutants had elevated expression relative to the control, consistent with the helix regulating endocytosis, and led to changes in CTxB and Lck staining (**Figure S4A-S4D**)(Sleckman et al., 1992).

Normalized sucrose data showed that the percent of CD4 localized to the DRMs and DSMs was not different for the H and HC mutants relative to the WT (**Fig 4A**). The HC mutant greatly reduced the normalized CTxB signal in the DRMs, while both H and HC reduced the signal in the DSMs (**Figure 4B**). They also reduced the normalized Lck signal associated with CD4 in the DRM and DSM relative to the wild-type (**Figure 4C**). These data indicate that mutating the helix alone relieves association with Lck.

**Figure 4.**
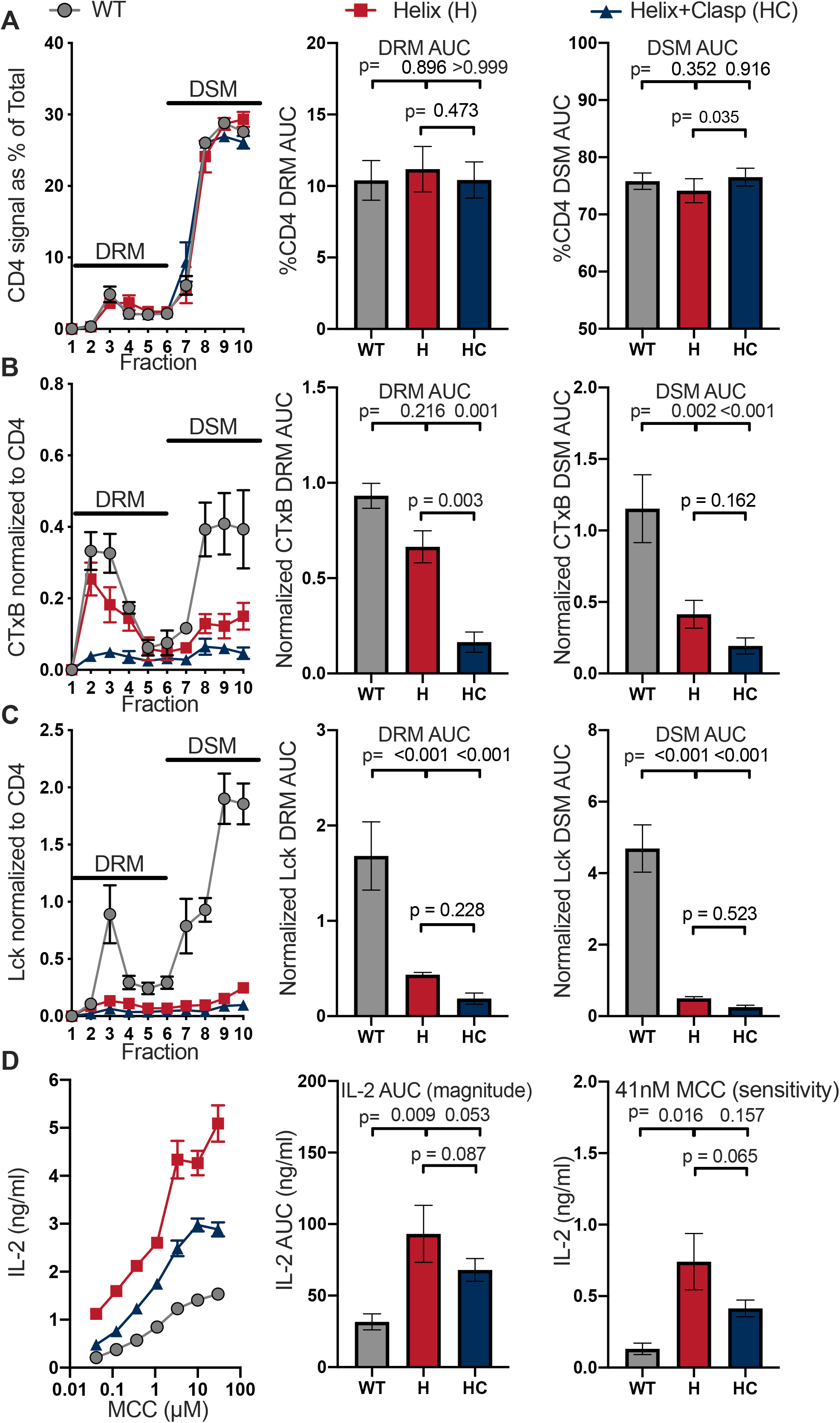
The intracellular helix attenuates response magnitude and sensitivity. (A) CD4 signal normalized as a percent of the total is shown for each sucrose gradient fraction (left) along with AUC analysis for the DRM (center) and DSM fractions (right). (B) Cholera toxin B (CTxB) signal normalized to CD4 signal detected is shown for each sucrose fraction (left) along with the AUC analysis for the DRM (center) and DSM fractions (right). (C) Lck signal normalized to CD4 signal is shown for each sucrose fraction (left) along with the AUC analysis for the DRM (center) and DSM fractions (right). (D) IL-2 dose response to MCC peptide (left), AUC analysis as a measure of the response magnitude (center), and the average response to a low dose (41nM) of MCC as a measure of sensitivity (right) are shown. For (A-D) the data are presented as in Figure 2. One-way ANOVA was performed with a Dunnett’s posttest for comparisons with WT samples and a Sidak’s posttest for comparisons between selected samples. See also Figure S4.

Given that the H and HC mutants failed to associate with Lck we expected both mutants to significantly impair pMHCII-responses; yet, we found that both the H and HC mutants increased the magnitude of the response across the peptide titration as well as the sensitivity to the lowest peptide dose (**Figure 4D**). Also, consistent with prior reports that the helix plays a role in endocytosis, the H and HC mutants did not show reduced surface expression after 16hrs in these functional assays (**Figure S4E**). These data show that without the helix, CD4-Lck association is not necessary for CD4 to enhance pMHCII responses.

### The IKRLL interacts with Lck and attenuates pMHCII responses

We next asked if specific residues within the helix were responsible for the H mutant phenotype. Accordingly, we mutated the dileucine repeat (L437A+L438A = LL) within the IKRLL motif because I/LXXLL motifs and dileucine repeats are known protein interaction sites that could mediate interactions with a variety of partners; likewise, we mutated the serine residues (S432A.S439A = SS) which could play a similar role upon (de)phosphorylation. The SS mutant had slightly lower expression than the WT, and both mutants showed variations in raw CTxB and Lck signal (**Figure S5A-S5D**). The normalized sucrose gradient data revealed minor variations in the percent CD4 signal in the DRMs and DSMs for the LL mutant and SS mutants relative to the WT (**Figure 5A**). Likewise, within the DRM and DSMs there were no remarkable difference in CTxB signal between the WT and mutants (**Figure 5B**). Finally, the LL mutant showed reduced association with Lck in the DRMs and DSMs relative to the WT, which is consistent with the IKRLL motif interacting with Lck (**Figure 5C**) (Kim et al., 2003; Sleckman et al., 1992). In contrast, the SS mutant slightly increased interactions with Lck in the DRM, compared to the WT, and greatly increased this interaction in DSM which is consistent with regulating CD4-Lck interactions (Sleckman et al., 1992). These data show that residues within the helix differentially regulate CD4 association with Lck.

**Figure 5.**
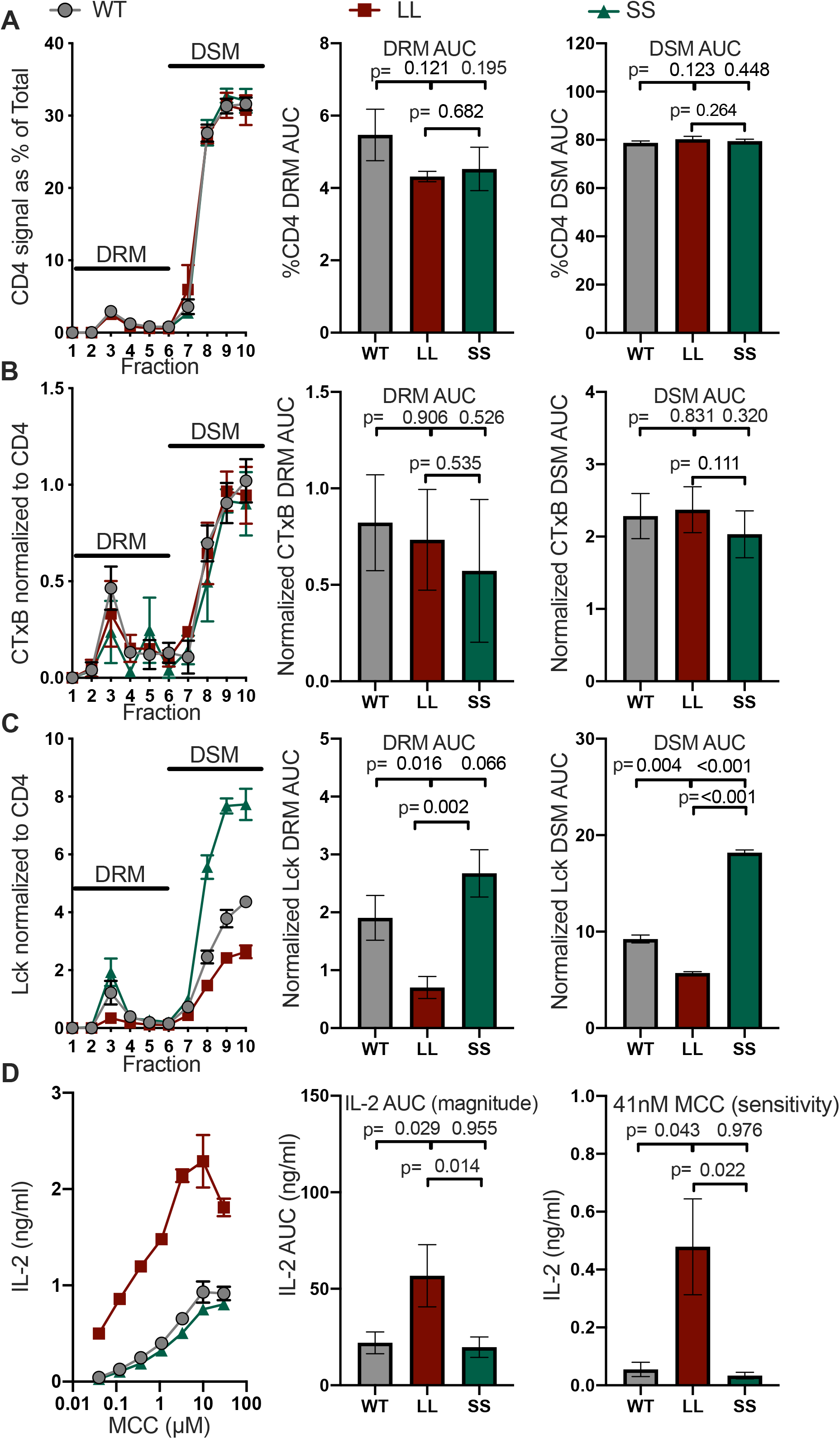
The IKRLL motif mediates the inhibitory function of the helix. (A) CD4 signal normalized as a percent of the total is shown for each sucrose gradient fraction (left) along with AUC analysis for the DRM (center) and DSM fractions (right). (B) Cholera toxin B (CTxB) signal normalized to CD4 signal detected is shown for each sucrose fraction (left) along with the AUC analysis for the DRM (center) and DSM fractions (right). (C) Lck signal normalized to CD4 signal is shown for each sucrose fraction (left) along with the AUC analysis for the DRM (center) and DSM fractions (right). (D) IL-2 dose response to MCC peptide (left), AUC analysis as a measure of the response magnitude (center), and the average response to a low dose (41nM) of MCC as a measure of sensitivity (right) are shown. For (A-D) the data are presented as in Figure 2. One-way ANOVA was performed with a Dunnett’s posttest for comparisons with WT samples and a Sidak’s posttest for comparisons between selected samples. See also Figure S5.

If CD4 association with Lck is the chief determinant for pMHCII-responses then the LL mutant should reduce IL-2 production by our 58α^-^β^-^ cells in response to cognate peptide while the SS mutant should enhance responses. However, we observed that the LL mutation greatly increased both the magnitude of the response as well as the sensitivity to a low peptide dose, while the SS did not differ from the WT (**Figure 5D**). Both mutations slightly impaired pMHCII-induced endocytosis of CD4, consistent with their previously repeated role in regulating this function (**Figure S5E**). Taken together with the H and HC mutant phenotypes, we interpret the sum of the data to indicate that the dominant function of the helix is to inhibit CD4’s contribution to pMHCII responses via the IKRLL motif.

### Coevolution of motifs in the ECD and ICD

Because a number of the motifs considered above co-arose in mammals or eutherians and, for the GGXXG and CV+C motifs, showed evidence of coordinated function, we were motivated to explore if residues in our regions of interest showed evidence of covariation. Constraints on protein function can lead to correlated mutations between residues in a protein that provide further evidence of their functional importance and can highlight networks of functional residues within a protein (Lockless and Ranganathan, 1999). We therefore used MISTIC2 to calculate the covariation between residues of CD4 (Colell et al., 2018; Kowarsch et al., 2010). MISTIC2 quantifies correlations using mutual information as a measure for how much information one random variable provides about another, allowing for detection of coevolutionary relationships between residues that are spatially distant and not just those that are in close proximity.

By analyzing the full data set we identified a network of interactions involving residues within the helix-turn. Specifically, five pairs of coevolving residues were found within the ICD helix-turn region (S432 – I434; S432 – S439; I434 – K441; L437 – S439; L437 – K442), which may be relevant to the structure of this region, its function, or both (**Figure 6**). Interestingly, G402 covaries with L438, suggesting a functional interaction between the GGXXG and IKRLL motifs as well as a covariation network between the TMD and ICD. Furthermore, H423 coevolved with I434 and S439, and R426 coevolved with S439. Prior structural data suggests that the HXXR is unstructured and not physically contacting the helix, suggesting instead a functional interplay between the HxxR motif and the helix (Kim et al., 2003). Remarkably, we also found that P228 and P281 of the nonpolar patch in the D3 domain of the ECD shows strong covariation with residues in the ICD helix. Specifically, P228 coevolved with S432, I434, S439, and K441, while P281 coevolved with S432, L438, and K442, providing evidence for the evolution of intracellular regulation of an extracellular event. These data led us to postulate that when the nonpolar patch arose, providing a mammalian CD4 ancestor with extracellular interactions that enhance pMHCII responses, the increased signaling was advantageous under some circumstances but detrimental in others; the conditions were thus set for the evolution of a network of regulatory motifs that allowed for finetuning of CD4’s role in the relay of pMHCII-specific information from the outside to the inside of a CD4^+^ T cell.

**Figure 6.**
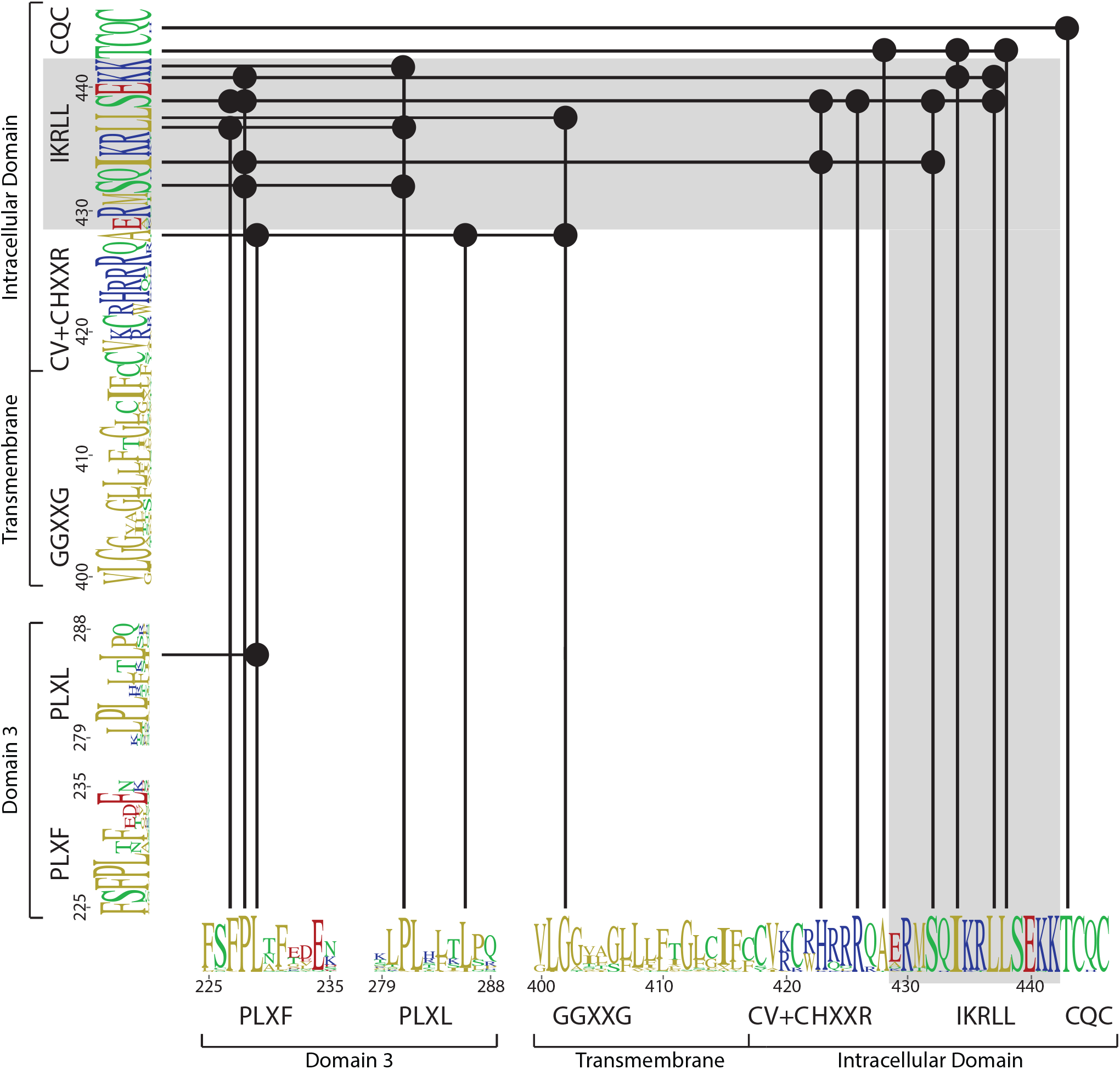
Covariation matrix. Covarying residues were calculated using MISTIC2. Residues that covary are indicated with a black dot and connected with a solid line. Motifs identified in this study are indicated. The logo plot represents eutherian sequences. The complete MISTIC2 results matrix is available on Dryad (https://doi.org/10.5061/dryad.59zw3r26z).

### Evidence for a functional interplay between coevolving motifs

To evaluate how the coevolution of these motifs impact pMHCII responses we assessed IL-2 responses by 58α^-^β^-^ cells expressing CD4 molecules that have both an enhancing motif and the inhibitor IKRLL motif mutated. We wanted to know if one motif would dominate over the other, or if we would observe an interplay between the motifs.

We combined an ELXE mutant (P228E+F231E) of the PLXF motif in the D3 domain, which we reported to reduce 58α^-^β^-^ IL-2 responses, with the LL mutant to assess if the helix inhibits contributions from the nonpolar patch (**Figure S6A**) (Glassman et al., 2018). We also combined a previously described GKGVLIR to GDGDSDS mutant in the D1 domain (CD4^Δbind^), which kills CD4 binding to pMHCII and pMHCII responses, to confirm that the LL mutant phenotype is dependent CD4-pMHCII interactions (Glassman et al., 2016; Parrish et al., 2015). We found that the LL + ELXE double mutant yielded similar results to the WT with regards to the magnitude and sensitivity of the response to MCC, while the LL + Δbind double mutant completely impaired responses (**Figure 7A and 7B**). Likewise, combining the TMD and Palm mutants with the dileucine mutant (TP+LL) reduced the magnitude of IL-2 production across a MCC titration, and at the lowest peptide dose, relative to the LL mutant such that the response was indistinguishable from the WT (**Figure 7A, 7B, and S6C**). Similarly, combining the R426A or Clasp mutants with the LL mutation generated similar responses to the WT.

**Figure 7.**
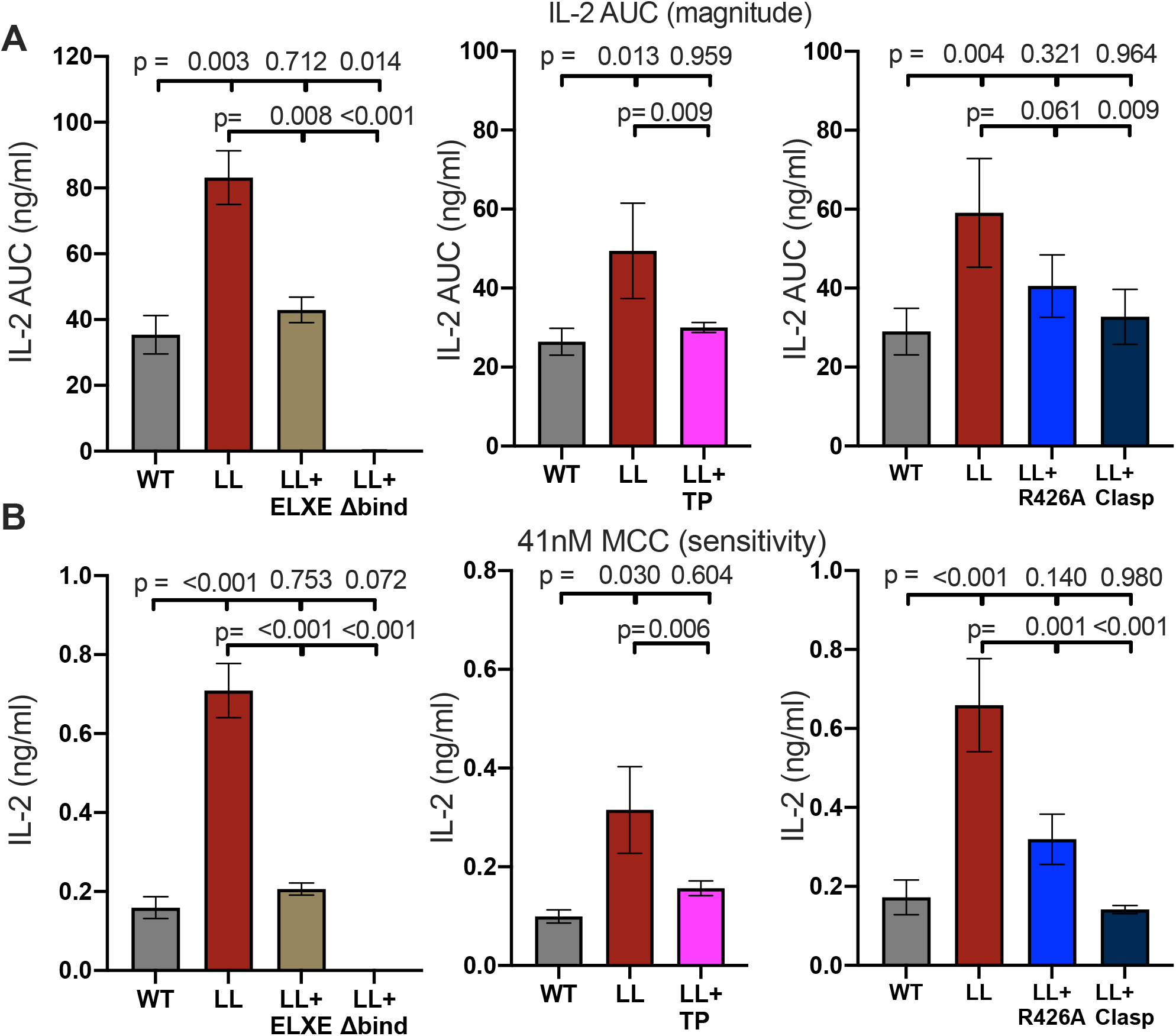
Coevolving motifs in the ECD and ICD functionally interact. (A) AUC analysis of IL-2 dose responses to MCC peptide are shown as a measure of the response magnitude for the indicated samples. (B) The average IL-2 response to a low dose (41nM) of MCC is shown as a measure of sensitivity for the indicated samples. For (A and B) the magnitude and sensitivity data represent the mean +/- SEM of three independent experiments (biological replicates) for which triplicate measurements were performed (technical replicates). One-way ANOVA was performed with a Dunnett’s posttest for comparisons with WT samples, and a Sidak’s posttest for comparisons between selected samples. Individual graphs indicate experiments that were performed with cell lines generated at the same time. See also Figure S6.

These data indicate that the LL mutation does not enhance pMHCII responses in the absence of CD4-pMHCII interactions. Rather, the ICD helix can counter-balance the impact of ECD, TMD, and ICD enhancing motifs. Our results with the LL + Clasp double mutant are also consistent with our HC mutant phenotype (**Figure 4D**), supporting the idea that CD4-Lck interactions are not necessary for pMHCII responses in the absence of the inhibitory function of the IKRLL motif. Together, the data provide evidence for a functional interplay between motifs that are coevolving with the inhibitory helix in the ICD to regulate events occurring outside and inside the cell.

## DISCUSSION

Biology has been experimenting with the form and function of CD4 *in vivo* for ∼435 million years in a variety of jawed vertebrates. The results of these experiments, as deconvolved by our computational reconstruction of CD4 evolution, thus provide insights into the residues or motifs that are essential for CD4 to confer fitness to jawed vertebrates within a particular clade or species in a way that cannot be readily replicated through reductionist studies of a single species or genetically engineered model thereof. Coupling our computational results with biochemical and functional analyses then provides us with mechanistic insights into why these residues or motifs are important for fitness. Together, our multidisciplinary approach provides evidence that eutherian CD4 evolved an epistatic network of function-regulating motifs wherein several of the enhancing motifs, including the nonpolar patch in the ECD, coevolved with an inhibitory helix in the ICD. The results provide insights into the evolution and function of the multi-subunit receptor system that relays pMHCII-specific information from the outside to the inside of CD4^+^ T cells.

The data presented here advance our understanding of eutherian CD4 by showing that the TMD and ICD evolved motifs that influence where Lck-associated CD4 molecules localize, and how well they drive pMHCII responses. As in prior work, we found that the CQC and CV+C motifs individually influence CD4 localization to DRMs (Fragoso et al., 2003). We extend this knowledge by showing that the CQC clasp, but not the CV+C motif alone, influences if CD4 molecules within DRMs inhabit GM1-containing rafts. Furthermore, we show that together the GGXXG and CV+C motifs localize Lck-associated CD4 molecules to DRMs and help localize CD4 molecules within DRMs to GM1-containing raft. We also extend prior insights about the HXXR motif by showing that it minimizes accumulation of Lck-associated CD4 molecules to DSMs, enhances CD4 expression, and enhances pMHCII responses (Popik and Alce, 2004). Similar results are presented for the HRΦQK motif. One interpretation of these data, which fits within the conceptual framework for the role of membrane rafts in cell signaling, is that eutherian CD4 is under selective pressure to regulate the membrane domain in which CD4 interacts with Lck because localization to membrane rafts, not simply interacting with Lck, is a key determinant for pMHCII responses. But this interpretation is only accurate when the IKRLL motif of the intracellular helix is intact, as without this motif CD4-Lck interactions are not necessary. We therefore conclude that the inhibitory function of the helix counterbalances the activity of the enhancing motifs to both prevent those CD4 molecules that are not associated with Lck from contributing to signaling and enforce a requirement that CD4-Lck interactions occur in membrane rafts for optimal pMHCII responses.

Our results also offer clues as to why and how eutherian CD4 evolved such elaborate control over CD4 activity. We find that the residue equivalent to P281 is present in all eutherian CD4 molecules but absent in the other amniotes we analyzed. This residue is a constituent of the nonpolar patch in the D3 domain of CD4 that contributes to the formation and stability of TCR-CD3-pMHCII-CD4 assemblies. We posit that the acquisition of P281 may have been a large-effect mutation, as described for other proteins, that conferred upon CD4 enhanced function and set the conditions for the rapid acquisition of additional mutations to balance the impact of the nonpolar patch (Whitney et al., 2016). A *de novo* insertion(s) between the TMD and the clasp of the amniote MRCA ICD (node 1), which likely explains the size difference between the fish and amniote CD4 ICD, would provide evolution with a substrate to explore novel sequences and add function to a previously non-functional region through the selection of beneficial mutations (Davey et al., 2015; Van Roey et al., 2012). In this scenario, individual motifs that impact the functional success of one another become dependent upon each other, as the fitness cost to change any single motif would be too large. This scenario is supported by the conservation of motifs we observe with residues under purifying selection, the covariation network we identified between the nonpolar patch in the ECD with the ICD helix, and our experimental analysis. It is also in line with the model of residue entrenchment described elsewhere (Shah et al., 2015).

The outcome for eutherian CD4 is the accumulation of coevolving motifs with switch-like capabilities to finetune the relay of pMHCII-specific information across the cell membrane. For example, the function of the CV+C motif is likely to be dynamic as palmitoylation is reversible (Ladygina et al., 2011). Furthermore, GXXXG motifs mediate interactions with other proteins, or with cholesterol which can modulate protein-protein interactions or induce allosteric changes (Song et al., 2014; Teese and Langosch, 2015). As cholesterol concentrations differ in distinct membrane domains, and can change upon TCR signaling, the GGXXG motif in eutherian CD4 may assume different functions depending on the membrane domain in which it is located or the activation or differentiation state of the T cell (Fessler, 2016). Of note, palmitate can interact with protein-bound cholesterol (PDB 4IB4), making it plausible that coupling of the palmitoylation state of the CV+C motif with a cholesterol-bound state of the GGXXG motif could further influence CD4 membrane domain localization and function. The HXXR and HRΦQK motifs are also likely to have dynamic functions as both are rich in basic residues that could form reversible electrostatic interactions with acidic headgroups of particular lipid species, serve as docking sites for low-affinity interactions with proteins or second messenger ligands, or both (Aivazian and Stern, 2000; Davey et al., 2015). For example, HXXR motifs are common in the P2Y family of GPCRs and related cysteinyl-leukotriene receptors, where they mediate ligand binding (Parravicini et al., 2010). Moreover, a RXXK motif in Slp-76 has been described as a nonproline-based binding site for the SH3 domains of Gads and Grb2; the histidine residue of the HRΦQK motif, which is under purifying selection, would not favor interactions with Gads and Grb2 according to that study, but it may define binding specificity for other SH3-domain bearing interaction partners (Berry et al., 2002). Of note, changes in membrane proximal cytoplasmic pH upon T cell activation could alter the charge of the histidine imidazole side-chain and favor particular interactions, perhaps to Gads or Grb2, under very specific conditions giving this motif switch like function. Finally, the helix appears to be a multifunctional hub, with basic residues that could mediate interactions with the membrane; serines that could be phosphorylated to repel the ICD from the membrane, stabilize interactions with other binding partners, or disrupt binding to Lck; and an IKRLL protein interaction motif that interacts with Lck but is likely to have other partners involved in its inhibitory function (Aivazian and Stern, 2000; Garron et al., 2008; Kelly et al., 2008; Sleckman et al., 1992; Yeung and Grinstein, 2007). Such an assembly of switch functions could thus provide tunability to CD4 function. Identifying interaction partners for these motifs, and deconvolving how they work in concert, will be the subject of future work.

In closing, the eutherian immune system achieves balance in its response to pMHCII not only through the interplay of helper and regulatory CD4^+^ T cells, or costimulatory and coinhibitory molecules, but also through finetuning CD4 function with enhancing and inhibitory motifs. Our findings are consistent with the idea that CD4 arose to enable more elaborate pMHCII responses than could be achieved otherwise (Venkatesh et al., 2014); indeed, it is interesting that the timing of key stepwise events in the evolution of *FoxP3*, or even helper and memory T cell functions, exhibit parallels with CD4 (Andersen et al., 2012; Lane et al., 2010). We expect that the coevolution of regulatory elements outside and inside of a cell, which to our knowledge is first evidenced here, will emerge as a theme for other transmembrane receptors tasked with relaying information to its intracellular signaling machinery. Finally, given that efforts to engineer synthetic receptors for therapeutic purposes are showing promise, most notably for chimeric antigen receptor (CAR)-T cell therapy, these results have translational value. The archetypal CAR design was conceived when it was understood that receptor bind ligands outside of a cell and generate signals inside the cell, but knowledge of the intermediate steps was lacking (Eshhar et al., 1993). Problems with sensitivity and side effects now suggest that the absence of mechanisms to mediate or regulate the relay of information across the membrane have a fitness cost for this form of CAR (Labanieh et al., 2018). We therefore think that biomimetic designs, incorporating strategies refined over ∼435 million years of iterative testing in a variety of vertebrates, will ultimately lead to more sensitive and reliable synthetic receptors (Kobayashi et al., 2020). Such biomimetic engineering will require a doubling down on basic research efforts to elucidate the evolutionary blueprint for key immune receptors.

## MATERIALS AND METHODS

### Evolutionary analyses

Available CD4 orthologs were downloaded from GenBank. The criteria used for including orthologous CD4 sequences in our analyses were that they have a domain structure consisting of four extracellular Ig domains followed by a TMD and a C-terminal ICD. Sequences that were shorter, contained frameshift mutations, etc…, were excluded from the analysis. Teleost fish were considered to be the oldest living species that contain a CD4 molecule in this study as a CD4 ortholog was not identified in cartilaginous fish (Venkatesh et al., 2014). The final dataset contained 99 unique CD4 orthologs, ranging from teleost fish to human. These (putative) coding sequences were translated to amino-acids and aligned using MAFFT (Katoh et al., 2002). For codon-based analyses, the aligned amino acid sequences were back translated to nucleotides to maintain codons. The multiple sequence alignments were further processed to remove all insertions (indels) relative to the mouse CD4 sequence (NM_013488.3) to maintain consistent numbering of sites. The 5’ and 3’ regions of the CD4 molecules were not consistently aligned due to different start codon usage or extensions of the ICD, respectively. The alignment was edited to start at the codon (AAG) coding for K48 within mouse CD4. The alignment that includes all 99 CD4 sequences ends at the last cysteine residue that makes up the CQC clasp. For the mammalian only dataset, the alignment terminates at the mouse CD4 stop codon.

FastTree was used to estimate maximum likelihood trees (Price et al., 2010). For amino acid-based trees, the Jones-Taylor-Thornton (JTT) model of evolution was selected, while the General Time-Reversible (GTR) model was used for nucleotide trees (Jones et al., 1992; Waddell and Steel, 1997). In all conditions, a discrete gamma model with 20 rate categories was used. A reduced representation tree (based on the amino acid alignment) is shown in Figure 1A. We used the same JTT model to estimate the marginal reconstructions of nodes indicated in Figure 1A. Phylogenetic trees and logo plots were visualized in Geneious and further edited using Adobe Illustrator. Ancestral sequences were estimated using GRASP (Foley et al., 2020).

Codon-based analysis of selection was performed using the hypothesis testing fixed effects likelihood (FEL) model as implemented within the phylogenies (HyPhy) package (version 2.5.14) (MP) (Kosakovsky Pond et al., 2020; Pond et al., 2005; Pond and Frost, 2005; Weaver et al., 2018). The back-translated codon-based alignments described above were used for these analyses. FEL uses likelihood ratio tests (LRTs) to assess a better fit of codons that allowed selection (*P* < 0.1). When calculating values for all CD4 orthologs included in the initial phylogenetic analysis we analyzed and identified the sequences within the mammalian clade as the foreground branches on which to test for evolutionary selection in order to maximize statistical power.

Covariation between protein residues was calculated using the MISTIC2 server. We calculated 4 different covariation methods (MIp, mFDCA, plmDCA, and gaussianDCA)(Colell et al., 2018). Protein conservation scores were calculated based on the protein alignment using the ConSurf Server (Ashkenazy et al., 2016; Ashkenazy et al., 2010). ConSurf conservation scores are normalized, so that the average score for all residues is zero, with a standard deviation of one. The lower the score, the more conserved the protein position. For the purpose of this study, residues were considered to covary if the MI was larger than 4 and both residues had a ConSurf conservation score lower than -0.5. Also, pairs with an MI larger than 8 were considered to covary if the conservation score was below -0.3. Using these criteria, we selected 0.5% of all possible pairs as recommended (Buslje et al., 2009; Colell et al., 2018). Raw data, including alignments and phylogenetic trees, associated with figures 1, S1, and 6 are available on Dryad (https://doi.org/10.5061/dryad.59zw3r26z)..

### Cell lines

58α^-^β^-^ T cell hybridoma lines were generated by retroviral transduction and maintained in culture by standard techniques as previously described (Glassman et al., 2018; Letourneur and Malissen, 1989). In brief, one day after transduction the cells were cultured in 5µg/ml puromycin(Invivogen) and 5µg/ml zeocin(ThermoFisher Scientific) in RPMI 1640 (Gibco) supplemented with 5% FBS (Atlanta Biologicals or Omega Scientific), Penicillin-Streptomycin-Glutamine (Cytiva), and 50µM βME (ThermoFisher Scientific). The next day drug concentrations were increased to 10µg/ml puromycin (Invivogen) and 100µg/ml zeocin (ThermoFisher Scientific) in 10ml in a T25 flask. Aliquot of 1×10^7^ cells were frozen at days 5, 7, and 9. Cells were thawed from the day 5 freeze and cultured for 3 days in 10µg/ml puromycin and 100µg/ml zeocin, and maintained below 1×10^6^ cells/ml to use in the functional assays. Cells used in the functional assays were grown to 0.8×10^6^ cells/ml density and replicates of three functional assays were performed every other day. If cells exceeded 1×10^6^ cells/ml at any point in the process they were discarded as they lose reactivity at high cell densities and a new set of vials was thawed. Typically, two independent WT and mutant pairs were generated for any given mutant and tested for IL-2 to gain further confidence in a response phenotype. When cells lines are presented together in a graph, that indicates that the cell lines (WT and mutants sets) were generated at the same time from the same parental cell stock.

Given the number of mutant CD4 cell lines generated and handled in this study, the identity of the transduced CD4 gene was verified by PCR sequencing at the conclusion of three independent functional assays. 58α^-^β^-^ T cell hybridomas were lysed using DirectPCR Tail Lysis Buffer (Viagen Biotech) with proteinase K (SIGMA) for 2 hours at 65°C. Cells were then heated at 95°C for ten minutes. Cell debris was pelleted, and the supernatants were saved. CD4 was amplified by PCR using Q5 DNA Hot Start Polymerase (New England BioLabs) in 0.2µM dNTP, 0.2µM primer concentration, Q5 Reaction buffer, and water. CD4 was amplified using the following primers:

5’ primer: acggaattccgctcgagcgccaccatggtgcgagccatctctctcttagg

3’ primer: ctagcaagcttgtcgactcaagatcttcattagatgagattatggctcttctgc

Product were purified using SpinSmart Nucleic Acid Purification Columns (Thomas Scientific) and sent to Eton Bioscience for sequencing with the following 5’ CD4 primer: gtctctgaggagcagaag.

The I-E^k+^ M12 cells used as antigen presenting cells (APCs) were previously reported (Glassman et al., 2018). Cells were cultured in RPMI 1640 (Gibco) supplemented with 5% FBS (Atlanta Biologicals or Omega Scientific), Pencillin-Streptomycin-Glutamine (Cytiva), 50µM βME (ThermoFisher Scientific), and 5µg/mL Puromycin (Invivogen) and 50µg/ml Zeocin (ThermoFisher).

### Flow cytometry

Cell surface expression of CD4 and TCR-CD3 complexes were measured by flow cytometry. In brief, cells were stained for 30 minutes at 4°C in FACS buffer (PBS, 2% FBS, and 0.02% sodium azide) using anti-CD4 (clone GK1.5, eFluor 450 conjugate, ThermoFisher Scientific), anti-TCRα (anti-Vα11, clone RR8-1, APC conjugate, ThermoFisher Scientific), anti-CD3ε (145-2C11, ThermoFisher Scientific) and GFP was detected as a measure of the TCRβ-GFP subunit. Analysis was performed on a Canto II or LSRII (BD Biosciences) at the Flow Cytometry Shared Resource at the University of Arizona. Flow cytometry data were analyzed with FlowJo Version 9 software (Becton, Dickinson & Company).

### Functional assays

IL-2 production was measured to quantify pMHCII responses. 5×10^4^ transduced 58α^-^β^-^ T cell hybridomas were cocultured with 1X10^5^ transduced I-E^k+^ M12 cells in a 96 well round bottom plate in the presence of titrating amounts of MCC 88-103 peptide (purchased from 21^st^ Century Biochemicals at >95% purity) starting at 30µM MCC and a 1:3 titration (Glassman et al., 2018). The supernatants were collected and assayed for IL-2 concentration by ELISA after 16 hours of co-culture at 37°C. Anti-mouse IL-2 (clone JES6-1A12, BioLegend) antibody was used to capture IL-2 from the supernatants, and biotin anti-mouse IL-2 (clone JES6-5H4, BioLegend) antibody was used as the secondary antibody. Streptavidin-HRP (BioLegend) and TMB substrate (BioLegend) were also used.

To assess engagement-induced endocytosis, CD4 surface levels were measured by flow cytometry 16hrs after coculture with APCs and peptide as described above for IL-2 quantification. 96 well plates containing cells were washed with ice cold FACs buffer (PBS, 2%FBS, 0.02% sodium azide), transferred to ice, and Fc receptors were blocked with Fc block mAb clone 2.4G2 for 15 minutes at 4°C prior to surface staining for 30 minutes at 4°C with anti-CD4 (clone GK1.5 EF450, Invitrogen) and anti-Vβ3 TCR clone (clone KJ25, BD Pharmingen) antibodies. Cells were washed with FACS buffer prior to analysis on a LSRII (BD Biosciences) at the Flow Cytometry Shared Resource at the University of Arizona. Flow cytometry data were analyzed with FlowJo Version 10 software (Becton, Dickinson & Company). The average of the geometric mean of the TCR or CD4 signal was taken for the triplicate of the post 58α^-^β^-^ cells cocultured with M12 I-E^k+^ cells at 0µM MCC concentration. Each value of the raw gMFI of TCR or CD4 for cells cultured at 10µM MCC was subtracted from the average gMFI at 0µM. The values show the change of gMFI from 0µM to 10µM.

### Sucrose gradient analysis

Membrane fractionation by sucrose gradient was performed similarly to previously described methods (Hur et al., 2003; Parrish et al., 2016). For cell lysis, 6×10^7^ 58α^-^β^-^ T cell hybridomas were harvested and washed 2x using TNE buffer (Tris, NaCl, EDTA). Cells were lysed on ice in 1% Triton-X detergent in TNE in a total volume of 1mL for 10 minutes and then dounce homogenized 10x. The homogenized lysate was transferred to 14 x 95mm Ultraclear Ultra Centrifuge tubes (Beckman). The dounce homogenizer was rinsed with 1.6mL of the 1% lysis buffer, which was then was added to the Ultracentrifuge tube. 2.5mL of 80% sucrose was added to the centrifuge tube with lysate, and mixed well. Gently, 5mL of 30% sucrose was added to the centrifuge tubes, creating a 30% sucrose layer above the ∼40% sucrose/lysate mixture. Then, 3mL of 5% sucrose was added gently to the centrifuge tube, creating another layer. The centrifuge tubes were spun 18 hours at 4°C in a SW40Ti rotor at 36,000rpm.

Analysis of membrane fractions was performed via flow-based fluorophore-linked immunosorbent assay (FFLISA) as previously described (Parrish et al., 2016). In brief, 88µL of Streptavidin Microspheres 6.0µm (Polysciences) were coated overnight at 4°C with 8µg of biotinylated anti-CD4 antibody (clone RM4-4, BioLegend). Prior to immunoprecipitation (IP), beads were washed with 10mL of FACS buffer (1x PBS, 2%FBS, 0.02% sodium azide) and resuspended in 3.5mL of 0.1% Triton-X 100 lysis wash buffer in TNE. For each cell line lysed, 10 FACS tubes were prepared with 50uL of the washed RM4-4 coated beads. Upon completion of the spin, 500µL was carefully taken off the top of the centrifuge tubes and discarded. Following this, 1mL was extracted from the top of the tube, carefully as to not disrupt the gradient, and added to a FACS tube with coated beads and capped. This was repeated for 10 individual fractions in separate FACS tubes. Following the extraction, lysates were incubated with the beads for 90 minutes, inverting the tubes to mix every 15 minutes.

Following the immunoprecipitation, FACS tubes were washed 3x using 0.1% Triton-X lysis wash buffer in TNE. Tubes were then stained using 1µL anti-CD4 (APC conjugate; clone GK1.5, BioLegend), 1.5µL anti-Lck (PE conjugate, clone 3A5, Santa Cruz Biotechnology), and 1µL CTxB (AF 488 conjugate, ThermoFisher Scientific, resuspended as per manufacturers instructions) for 45 minutes at 4°C. Following the stain, tubes were washed using 0.1% Triton-X lysis wash buffer in TNE. Analysis of beads were performed on a LSRII (BD Biosciences) at the Flow Cytometry Shared Resource at the University of Arizona. 10^4^ events were collected per sample. Flow cytometry data were analyzed with FlowJo Version 10 software (Becton, Dickinson & Company).

For FFLISA analysis, raw gMFI values for fraction 1 were subtracted from the rest of the fractions to account for background, such that the gMFI of fraction 1 is 0. To normalize the data, the percentage of CD4 within any given fraction (fx) relative to the total CD4 gMFI (CD4 signal % of total) was calculated by dividing the gMFI signal in a given fraction (fx) by the sum of the total CD4 gMFI signal [sum(f1:f10)CD4 gMFI] and multiplying by 100 [e.g. fx % of total = fx CD4 gMFI / Sum(f1:f10)CD4 gMFI x 100]. To normalize the CTxB and Lck signal in any given fraction relative to the CD4 signal in that same fraction (CTxB or Lck normalized to CD4) the gMFI of CTxB or Lck in fx was divided by the CD4 gMFI of that fx and then multiplied by the percentage of CD4 within fx (e.g. Normalized fx Lck = fx Lck gMFI/fx CD4 gMFI x fx CD4% of total CD4 gMFI). Area under the curve (AUC) analysis was performed with GraphPad Prism 9 for fractions 1-6 to determine the AUC for the detergent resistant membrane (DRM) domains due to their floating phenotypes, and for fractions 6-10 to determine the AUC for the detergent soluble membrane (DSM) domains.

### Statistical Analysis

Statistical analyses of sucrose gradient and functional assays were performed with GraphPad Prism 9 software as indicated in the figure legends. For each functional assay (IL-2 production and CD4 endocytosis), each individual experiment (biological replicate) was performed with triplicate analysis (technical replicates) and each experiment was repeated three times (3 biological replicates). For sucrose gradient analysis, 10^4^ beads were collected by flow cytometry in each experiment (technical replicates) and each experiment was performed three times (biological replicates). Three biological replicates were chosen for each analysis as per convention, and no power calculations were determined. One-Way ANOVA were performed with a Dunnett’s posttest when all mutants tested in an experiment were compared to a control sample (e.g. WT). Sidak’s posttest were applied when comparing between two specific samples. These posttests were chosen based on Prism recommendations.

### Constructs

The following sequence for 5cc7α was subcloned into the pZ4 zeocin-resistance MSCV vector (MCS-IRES-Zeo) resistance through 5’XhoI and 3’EcoRI:

cctcgagcgccaccatggagaggaacctgggagctgtgctggggattctgtgggtgcagatttgctgggtgagaggagat
caggtggagcagagtccttcagccctgagcctccacgagggaaccggttctgctctgagatgcaattttactaccaccatg
agggctgtgcagtggttccgaaagaattccaggggcagcctcatcaatctgttctacttggcttcaggaacaaaggagaat
gggaggctaaagtcagcatttgattctaaggagcgctacagcaccctgcacatcagggatgcccagctggaggactcag
gcacttacttctgtgctgctgaggcttccaataccaacaaagtcgtctttggaacagggaccagattacaagtattaccaaa
catccagaacccagaacctgctgtgtaccagttaaaagatcctcggtctcaggacagcaccctctgcctgttcaccgacttt
gactcccaaatcaatgtgccgaaaaccatggaatctggaacgttcatcactgacaaaactgtgctggacatgaaagctat
ggattccaagagcaatggggccattgcctggagcaaccagacaagcttcacctgccaagatatcttcaaagagaccaa
cgccacctaccccagttcagacgttccctgtgatgccacgttgaccgagaaaagctttgaaacagatatgaacctaaacttt
caaaacctgtcagttatgggactccgaatcctcctgctgaaagtagcgggatttaacctgctcatgacgctgaggctgtggt
ccagtgcggccgcatgataaagatctggatcctgactgagtgagaattc

The following sequence for 5cc7βG was subcloned into the pP2 puromycin-resistance MSCV vector (MCS-IRES-puro) resistance through 5’XhoI and 3’BamHI:

actcgagcgccaccatggctacaaggctcctctgttacacagtactttgtctcctgggtgcaagaattttgaattcaaaagtc attcagactccaagatatctggtgaaagggcaaggacaaaaagcaaagatgaggtgtatccctgaaaagggacatcca gttgtattctggtatcaacaaaataagaacaatgagtttaaatttttgattaactttcagaatcaagaagttcttcagcaaatag acatgactgaaaaacgattctctgctgagtgtccttcaaactcaccttgcagcctagaaattcagtcctctgaggcaggaga ctcagcactgtacctctgtgccagcagtctgaacaatgcaaactccgactacaccttcggctcagggaccaggcttttggta atagaggatctgagaaatgtgactccacccaaggtctccttgtttgagccatcaaaagcagagattgcaaacaaacaaa aggctaccctcgtgtgcttggccaggggcttcttccctgaccacgtggagctgagctggtgggtgaatggcaaggaggtcc acagtggggtcagcacggaccctcaggcctacaaggagagcaattatagccactgcctgagcagccgcctgagggtct ctgctaccttctggcacaatcctcgcaaccacttccgctgccaagtgcagttccatgggctttcagaggaggacaagtggc cagagggctcacccaaacctgtcacacagaacatcagtgcagaggcctggggccgagcagactgtgggattacctca gcatcctatcaacaaggggtcttgtctgccaccatcctctatgagatcctgctagggaaagccaccctgtatgctgtgcttgtc agtacactggtggtgatggctatggtcaaaagaaagaattccgcggccgcaggtggaggcggatcaggtggcggtgga agtggaggtggtggatctatggtgagcaagggcgaggagctgttcaccggggtggtgcccatcctggtcgagctggacg gcgacgtaaacggccacaagttcagcgtgtccggcgagggcgagggcgatgccacctacggcaagctgaccctgaa gttcatctgcaccaccggcaagctgcccgtgccctggcccaccctcgtgaccaccctgacctacggcgtgcagtgcttcag ccgctaccccgaccacatgaagcagcacgacttcttcaagtccgccatgcccgaaggctacgtccaggagcgcaccat cttcttcaaggacgacggcaactacaagacccgcgccgaggtgaagttcgagggcgacaccctggtgaaccgcatcga gctgaagggcatcgacttcaaggaggacggcaacatcctggggcacaagctggagtacaactacaacagccacaac gtctatatcatggccgacaagcagaagaacggcatcaaggtgaacttcaagatccgccacaacatcgaggacggcag cgtgcagctcgccgaccactaccagcagaacacccccatcggcgacggccccgtgctgctgcccgacaaccactacct gagcacccagtccaagctgagcaaagaccccaacgagaagcgcgatcacatggtcctgctggagttcgtgaccgccg ccgggatcactctcggcatggacgagctgtacaagtaatgaggatcctga

Full-length CD3δ, ε, γ, and ζ were encoded on a poly-cistronic construct as previously described (Holst et al., 2008; Kuhns and Davis, 2007)

CD4 WT, CD4 TMD, CD4 Palm, CD4 Clasp, CD4 TP, CD4 TPC were cloned by conventional molecular biology techniques, including PCR-based mutagenesis where needed, into pUC18 via 5’EcoRI and 3’HindIII. After sequence verification they were subcloned into pP2 puromycin-resistance MSCV vector (MCS-IRES-puro) vectors via 5’XhoI and 3’BglII, midi-prepped (Qiagen), and sequence verified. The sequences for these constructs are as follows. Mutated codons are shown in bold uppercase letters, while the motif under interrogation is shown in bold and underlined (any non-mutated codons within the motif are bold lowercase letters and underlined):

#### CD4 WT

acggaattccgctcgagcgccaccatggtgcgagccatctctcttaggcgcttgctgctgctgctgctgcagctgtcacaac tcctagctgtcactcaagggaagacgctggtgctggggaaggaaggggaatcagcagaactgccctgcgagagttccc agaagaagatcacagtcttcacctggaagttctctgaccagaggaagattctggggcagcatggcaaaggtgtattaatta gaggaggttcgccttcgcagtttgatcgttttgattccaaaaaaggggcatgggagaaaggatcgtttcctctcatcatcaat aaacttaagatggaagactctcagacttatatctgtgagctggagaacaggaaagaggaggtggagttgtgggtgttcaa agtgaccttcagtccgggtaccagcctgttgcaagggcagagcctgaccctgaccttggatagcaactctaaggtctctaa ccccttgacagagtgcaaacacaaaaagggtaaagttgtcagtggttccaaagttctctccatgtccaacctaagggttca ggacagcgacttctggaactgcaccgtgaccctggaccagaaaaagaactggttcggcatgacactctcagtgctgggtt ttcagagcacagctatcacggcctataagagtgagggagagtcagcggagttctccttcccactcaactttgcagaggaa aacgggtggggagagctgatgtggaaggcagagaaggattctttcttccagccctggatctccttctccataaagaacaa agaggtgtccgtacaaaagtccaccaaagacctcaagctccagctgaaggaaacgctcccactcaccctcaagatacc ccaggtctcgcttcagtttgctggttctggcaacctgactctgactctggacaaagggacactgcatcaggaagtgaacctg gtggtgatgaaagtggctcagctcaacaatactttgacctgtgaggtgatgggacctacctctcccaagatgagactgacc ctgaagcaggagaaccaggaggccagggtctctgaggagcagaaagtagttcaagtggtggcccctgagacagggct gtggcagtgtctactgagtgaaggtgataaggtcaagatggactccaggatccaggttttatccagaggggtgaaccaga cagtgttcctggcttgcgtgctgggtggctccttcggctttctgggtttccttgggctctgcatcctctgctgtgtcaggtgccggc accaacagcgccaggcagcacgaatgtctcagatcaagaggctcctcagtgagaagaagacctgccagtgcccccac cggatgcagaagagccataatctcatctaatgaagatcttgagtcgacaagcttgctag

#### CD4 TMD

acggaattccgctcgagcgccaccatggtgcgagccatctctcttaggcgcttgctgctgctgctgctgcagctgtcacaactcctagctgtcactcaagggaagacgctggtgctggggaaggaaggggaatcagcagaactgccctgcgagagttcccagaagaagatcacagtcttcacctggaagttctctgaccagaggaagattctggggcagcatggcaaaggtgtattaattagaggaggttcgccttcgcagtttgatcgttttgattccaaaaaaggggcatgggagaaaggatcgtttcctctcatcatcaataaacttaagatggaagactctcagacttatatctgtgagctggagaacaggaaagaggaggtggagttgtgggtgttcaaagtgaccttcagtccgggtaccagcctgttgcaagggcagagcctgaccctgaccttggatagcaactctaaggtctctaaccccttgacagagtgcaaacacaaaaagggtaaagttgtcagtggttccaaagttctctccatgtccaacctaagggttcaggacagcgacttctggaactgcaccgtgaccctggaccagaaaaagaactggttcggcatgacactctcagtgctgggttttcagagcacagctatcacggcctataagagtgagggagagtcagcggagttctccttcccactcaactttgcagaggaaaacgggtggggagagctgatgtggaaggcagagaaggattctttcttccagccctggatctccttctccataaagaacaaagaggtgtccgtacaaaagtccaccaaagacctcaagctccagctgaaggaaacgctcccactcaccctcaagataccccaggtctcgcttcagtttgctggttctggcaacctgactctgactctggacaaagggacactgcatcaggaagtgaacctggtggtgatgaaagtggctcagctcaacaatactttgacctgtgaggtgatgggacctacctctcccaagatgagactgaccctgaagcaggagaaccaggaggccagggtctctgaggagcagaaagtagttcaagtggtggcccctgagacagggctgtggcagtgtctactgagtgaaggtgataaggtcaagatggactccaggatccaggttttatccagaggggtgaaccaga cagtgttcctggcttgcgtgctgggt**<u>GTGtccttcCTC</u>**tttctgggtttccttgggctctgcatcctctgctgtgtcaggtgccg gcaccaacagcgccaggcagcacgaatgtctcagatcaagaggctcctcagtgagaagaagacctgccagtgcccccaccggatgcagaagagccataatctcatctaatgaagatcttgagtcgacaagcttgctag

#### CD4 Palm

acggaattccgctcgagcgccaccatggtgcgagccatctctcttaggcgcttgctgctgctgctgctgcagctgtcacaactcctagctgtcactcaagggaagacgctggtgctggggaaggaaggggaatcagcagaactgccctgcgagagttcccagaagaagatcacagtcttcacctggaagttctctgaccagaggaagattctggggcagcatggcaaaggtgtattaattagaggaggttcgccttcgcagtttgatcgttttgattccaaaaaaggggcatgggagaaaggatcgtttcctctcatcatcaataaacttaagatggaagactctcagacttatatctgtgagctggagaacaggaaagaggaggtggagttgtgggtgttcaaagtgaccttcagtccgggtaccagcctgttgcaagggcagagcctgaccctgaccttggatagcaactctaaggtctctaaccccttgacagagtgcaaacacaaaaagggtaaagttgtcagtggttccaaagttctctccatgtccaacctaagggttcaggacagcgacttctggaactgcaccgtgaccctggaccagaaaaagaactggttcggcatgacactctcagtgctgggttttcagagcacagctatcacggcctataagagtgagggagagtcagcggagttctccttcccactcaactttgcagaggaaaacgggtggggagagctgatgtggaaggcagagaaggattctttcttccagccctggatctccttctccataaagaacaaagaggtgtccgtacaaaagtccaccaaagacctcaagctccagctgaaggaaacgctcccactcaccctcaagataccccaggtctcgcttcagtttgctggttctggcaacctgactctgactctggacaaagggacactgcatcaggaagtgaacctggtggtgatgaaagtggctcagctcaacaatactttgacctgtgaggtgatgggacctacctctcccaagatgagactgaccctgaagcaggagaaccaggaggccagggtctctgaggagcagaaagtagttcaagtggtggcccctgagacagggctgtggcagtgtctactgagtgaaggtgataaggtcaagatggactccaggatccaggttttatccagaggggtgaaccaga cagtgttcctggcttgcgtgctgggtggctccttcggctttctgggtttccttgggctctgcatcctctgc**<u>TCTgtcaggTCC</u>**c ggcaccaacagcgccaggcagcacgaatgtctcagatcaagaggctcctcagtgagaagaagacctgccagtgcccccaccggatgcagaagagccataatctcatctaatgaagatcttgagtcgacaagcttgctag

#### CD4 Clasp

acggaattccgctcgagcgccaccatggtgcgagccatctctcttaggcgcttgctgctgctgctgctgcagctgtcacaactcctagctgtcactcaagggaagacgctggtgctggggaaggaaggggaatcagcagaactgccctgcgagagttcccagaagaagatcacagtcttcacctggaagttctctgaccagaggaagattctggggcagcatggcaaaggtgtattaattagaggaggttcgccttcgcagtttgatcgttttgattccaaaaaaggggcatgggagaaaggatcgtttcctctcatcatcaataaacttaagatggaagactctcagacttatatctgtgagctggagaacaggaaagaggaggtggagttgtgggtgttcaa agtgaccttcagtccgggtaccagcctgttgcaagggcagagcctgaccctgaccttggatagcaactctaaggtctctaaccccttgacagagtgcaaacacaaaaagggtaaagttgtcagtggttccaaagttctctccatgtccaacctaagggttcaggacagcgacttctggaactgcaccgtgaccctggaccagaaaaagaactggttcggcatgacactctcagtgctgggttttcagagcacagctatcacggcctataagagtgagggagagtcagcggagttctccttcccactcaactttgcagaggaaaacgggtggggagagctgatgtggaaggcagagaaggattctttcttccagccctggatctccttctccataaagaacaaagaggtgtccgtacaaaagtccaccaaagacctcaagctccagctgaaggaaacgctcccactcaccctcaagataccccaggtctcgcttcagtttgctggttctggcaacctgactctgactctggacaaagggacactgcatcaggaagtgaacctggtggtgatgaaagtggctcagctcaacaatactttgacctgtgaggtgatgggacctacctctcccaagatgagactgaccctgaagcaggagaaccaggaggccagggtctctgaggagcagaaagtagttcaagtggtggcccctgagacagggctgtggcagtgtctactgagtgaaggtgataaggtcaagatggactccaggatccaggttttatccagaggggtgaaccagacagtgttcctggcttgcgtgctgggtggctccttcggctttctgggtttccttgggctctgcatcctctgctgtgtcaggtgccggc accaacagcgccaggcagcacgaatgtctcagatcaagaggctcctcagtgagaagaagacc**<u>TCCcagTCC</u>**ccc caccggatgcagaagagccataatctcatctaatgaagatcttgagtcgacaagcttgctag

#### CD4 TP

acggaattccgctcgagcgccaccatggtgcgagccatctctcttaggcgcttgctgctgctgctgctgcagctgtcacaactcctagctgtcactcaagggaagacgctggtgctggggaaggaaggggaatcagcagaactgccctgcgagagttcccagaagaagatcacagtcttcacctggaagttctctgaccagaggaagattctggggcagcatggcaaaggtgtattaattagaggaggttcgccttcgcagtttgatcgttttgattccaaaaaaggggcatgggagaaaggatcgtttcctctcatcatcaataaacttaagatggaagactctcagacttatatctgtgagctggagaacaggaaagaggaggtggagttgtgggtgttcaaagtgaccttcagtccgggtaccagcctgttgcaagggcagagcctgaccctgaccttggatagcaactctaaggtctctaaccccttgacagagtgcaaacacaaaaagggtaaagttgtcagtggttccaaagttctctccatgtccaacctaagggttcaggacagcgacttctggaactgcaccgtgaccctggaccagaaaaagaactggttcggcatgacactctcagtgctgggttttcagagcacagctatcacggcctataagagtgagggagagtcagcggagttctccttcccactcaactttgcagaggaaaacgggtggggagagctgatgtggaaggcagagaaggattctttcttccagccctggatctccttctccataaagaacaaagaggtgtccgtacaaaagtccaccaaagacctcaagctccagctgaaggaaacgctcccactcaccctcaagataccccaggtctcgcttcagtttgctggttctggcaacctgactctgactctggacaaagggacactgcatcaggaagtgaacctggtggtgatgaaagtggctcagctcaacaatactttgacctgtgaggtgatgggacctacctctcccaagatgagactgaccctgaagcaggagaaccaggaggccagggtctctgaggagcagaaagtagttcaagtggtggcccctgagacagggctgtggcagtgtctactgagtgaaggtgataaggtcaagatggactccaggatccaggttttatccagaggggtgaaccaga cagtgttcctggcttgcgtgctgggt**<u>GTGtccttcCTC</u>**tttctgggtttccttgggctctgcatcctctgc**<u>TCTgtcaggTC C</u>**cggcaccaacagcgccaggcagcacgaatgtctcagatcaagaggctcctcagtgagaagaagacctgccagtgc ccccaccggatgcagaagagccataatctcatctaatgaagatcttgagtcgacaagcttgctag

#### CD4 TPC

acggaattccgctcgagcgccaccatggtgcgagccatctctcttaggcgcttgctgctgctgctgctgcagctgtcacaactcctagctgtcactcaagggaagacgctggtgctggggaaggaaggggaatcagcagaactgccctgcgagagttcccagaagaagatcacagtcttcacctggaagttctctgaccagaggaagattctggggcagcatggcaaaggtgtattaattagaggaggttcgccttcgcagtttgatcgttttgattccaaaaaaggggcatgggagaaaggatcgtttcctctcatcatcaataaacttaagatggaagactctcagacttatatctgtgagctggagaacaggaaagaggaggtggagttgtgggtgttcaaagtgaccttcagtccgggtaccagcctgttgcaagggcagagcctgaccctgaccttggatagcaactctaaggtctctaaccccttgacagagtgcaaacacaaaaagggtaaagttgtcagtggttccaaagttctctccatgtccaacctaagggttcaggacagcgacttctggaactgcaccgtgaccctggaccagaaaaagaactggttcggcatgacactctcagtgctgggttttcagagcacagctatcacggcctataagagtgagggagagtcagcggagttctccttcccactcaactttgcagaggaaaacgggtggggagagctgatgtggaaggcagagaaggattctttcttccagccctggatctccttctccataaagaacaaagaggtgtccgtacaaaagtccaccaaagacctcaagctccagctgaaggaaacgctcccactcaccctcaagataccccaggtctcgcttcagtttgctggttctggcaacctgactctgactctggacaaagggacactgcatcaggaagtgaacctg gtggtgatgaaagtggctcagctcaacaatactttgacctgtgaggtgatgggacctacctctcccaagatgagactgaccctgaagcaggagaaccaggaggccagggtctctgaggagcagaaagtagttcaagtggtggcccctgagacagggctgtggcagtgtctactgagtgaaggtgataaggtcaagatggactccaggatccaggttttatccagaggggtgaaccaga cagtgttcctggcttgcgtgctgggt**<u>GTGtccttcCTC</u>**tttctgggtttccttgggctctgcatcctctgc**<u>TCTgtcaggTC C</u>**cggcaccaacagcgccaggcagcacgaatgtctcagatcaagaggctcctcagtgagaagaagacc**<u>TCCcagT CC</u>**ccccaccggatgcagaagagccataatctcatctaatgaagatcttgagtcgacaagcttgctag

Gene blocks were purchased from Twist Biosciences that encode the required mutations in each of the following CD4 genes: R426A, R449A, K452A, Dbl, H, HC, LL, SS, LL+TP, LL+R426A, and LL+C. The gene segments were cloned into pUC18 containing CD4 WT whereby the gene segments flanked by BamHI and HindIII were replaced with the mutant geneblocks. The genes were sequence verified in pUC 18, subcloned into pP2 puromycin-resistance MSCV vector (MCS-IRES-puro) via 5’XhoI and 3’BglII, midi-prepped, and sequenced. The sequences for these constructs are listed below. Mutated codons are shown in bold uppercase letters, while the motif under interrogation is shown in bold and underlined (any non-mutated codons within the motif are bold lowercase letters and underlined):

#### CD4 R426A

acggaattccgctcgagcgccaccatggtgcgagccatctctcttaggcgcttgctgctgctgctgctgcagctgtcacaactcctagctgtcactcaagggaagacgctggtgctggggaaggaaggggaatcagcagaactgccctgcgagagttcccagaagaagatcacagtcttcacctggaagttctctgaccagaggaagattctggggcagcatggcaaaggtgtattaattagaggaggttcgccttcgcagtttgatcgttttgattccaaaaaaggggcatgggagaaaggatcgtttcctctcatcatcaataaacttaagatggaagactctcagacttatatctgtgagctggagaacaggaaagaggaggtggagttgtgggtgttcaaagtgaccttcagtccgggtaccagcctgttgcaagggcagagcctgaccctgaccttggatagcaactctaaggtctctaaccccttgacagagtgcaaacacaaaaagggtaaagttgtcagtggttccaaagttctctccatgtccaacctaagggttcaggacagcgacttctggaactgcaccgtgaccctggaccagaaaaagaactggttcggcatgacactctcagtgctgggttttcagagcacagctatcacggcctataagagtgagggagagtcagcggagttctccttcccactcaactttgcagaggaaaacgggtggggagagctgatgtggaaggcagagaaggattctttcttccagccctggatctccttctccataaagaacaaagaggtgtccgtacaaaagtccaccaaagacctcaagctccagctgaaggaaacgctcccactcaccctcaagataccccaggtctcgcttcagtttgctggttctggcaacctgactctgactctggacaaagggacactgcatcaggaagtgaacctggtggtgatgaaagtggctcagctcaacaatactttgacctgtgaggtgatgggacctacctctcccaagatgagactgaccctgaagcaggagaaccaggaggccagggtctctgaggagcagaaagtagttcaagtggtggcccctgagacagggctgtggcagtgtctactgagtgaaggtgataaggtcaagatggactccaggatccaggttttatccagaggggtgaaccaga cagtgttcctggcttgcgtgctgggtggctccttcggctttctgggtttccttgggctctgcatcctctgctgtgtcaggtgccgg**<u>c accaacagGCT</u>**caggcagcacgaatgtctcagatcaagaggctcctcagtgagaagaagacctgccagtgccccc accggatgcagaagagccataatctcatctgataaagatcttgagtcgacaagcttgctag

#### CD4 R449A

acggaattccgctcgagcgccaccatggtgcgagccatctctcttaggcgcttgctgctgctgctgctgcagctgtcacaactcctagctgtcactcaagggaagacgctggtgctggggaaggaaggggaatcagcagaactgccctgcgagagttcccagaagaagatcacagtcttcacctggaagttctctgaccagaggaagattctggggcagcatggcaaaggtgtattaattagaggaggttcgccttcgcagtttgatcgttttgattccaaaaaaggggcatgggagaaaggatcgtttcctctcatcatcaat aaacttaagatggaagactctcagacttatatctgtgagctggagaacaggaaagaggaggtggagttgtgggtgttcaaagtgaccttcagtccgggtaccagcctgttgcaagggcagagcctgaccctgaccttggatagcaactctaaggtctctaaccccttgacagagtgcaaacacaaaaagggtaaagttgtcagtggttccaaagttctctccatgtccaacctaagggttcaggacagcgacttctggaactgcaccgtgaccctggaccagaaaaagaactggttcggcatgacactctcagtgctgggttttcagagcacagctatcacggcctataagagtgagggagagtcagcggagttctccttcccactcaactttgcagaggaaaacgggtggggagagctgatgtggaaggcagagaaggattctttcttccagccctggatctccttctccataaagaacaaagaggtgtccgtacaaaagtccaccaaagacctcaagctccagctgaaggaaacgctcccactcaccctcaagataccccaggtctcgcttcagtttgctggttctggcaacctgactctgactctggacaaagggacactgcatcaggaagtgaacctggtggtgatgaaagtggctcagctcaacaatactttgacctgtgaggtgatgggacctacctctcccaagatgagactgaccctgaagcaggagaaccaggaggccagggtctctgaggagcagaaagtagttcaagtggtggcccctgagacagggctgtggcagtgtctactgagtgaaggtgataaggtcaagatggactccaggatccaggttttatccagaggggtgaaccagacagtgttcctggcttgcgtgctgggtggctccttcggctttctgggtttccttgggctctgcatcctctgctgtgtcaggtgccggc accaacagcgccaggcagcacgaatgtctcagatcaagaggctcctcagtgagaagaagacctgccagtgcccc**<u>cac GCTatgcagaag</u>**agccataatctcatctgataaagatcttgagtcgacaagcttgctag

#### CD4 K452A

acggaattccgctcgagcgccaccatggtgcgagccatctctcttaggcgcttgctgctgctgctgctgcagctgtcacaactcctagctgtcactcaagggaagacgctggtgctggggaaggaaggggaatcagcagaactgccctgcgagagttcccagaagaagatcacagtcttcacctggaagttctctgaccagaggaagattctggggcagcatggcaaaggtgtattaattagaggaggttcgccttcgcagtttgatcgttttgattccaaaaaaggggcatgggagaaaggatcgtttcctctcatcatcaataaacttaagatggaagactctcagacttatatctgtgagctggagaacaggaaagaggaggtggagttgtgggtgttcaaagtgaccttcagtccgggtaccagcctgttgcaagggcagagcctgaccctgaccttggatagcaactctaaggtctctaaccccttgacagagtgcaaacacaaaaagggtaaagttgtcagtggttccaaagttctctccatgtccaacctaagggttcaggacagcgacttctggaactgcaccgtgaccctggaccagaaaaagaactggttcggcatgacactctcagtgctgggttttcagagcacagctatcacggcctataagagtgagggagagtcagcggagttctccttcccactcaactttgcagaggaaaacgggtggggagagctgatgtggaaggcagagaaggattctttcttccagccctggatctccttctccataaagaacaaagaggtgtccgtacaaaagtccaccaaagacctcaagctccagctgaaggaaacgctcccactcaccctcaagataccccaggtctcgcttcagtttgctggttctggcaacctgactctgactctggacaaagggacactgcatcaggaagtgaacctggtggtgatgaaagtggctcagctcaacaatactttgacctgtgaggtgatgggacctacctctcccaagatgagactgaccctgaagcaggagaaccaggaggccagggtctctgaggagcagaaagtagttcaagtggtggcccctgagacagggctgtggcagtgtctactgagtgaaggtgataaggtcaagatggactccaggatccaggttttatccagaggggtgaaccagacagtgttcctggcttgcgtgctgggtggctccttcggctttctgggtttccttgggctctgcatcctctgctgtgtcaggtgccggc accaacagcgccaggcagcacgaatgtctcagatcaagaggctcctcagtgagaagaagacctgccagtgcccc**<u>cac cggatgcagGCA</u>**agccataatctcatctgataaagatcttgagtcgacaagcttgctag

#### CD4 Dbl

acggaattccgctcgagcgccaccatggtgcgagccatctctcttaggcgcttgctgctgctgctgctgcagctgtcacaactcctagctgtcactcaagggaagacgctggtgctggggaaggaaggggaatcagcagaactgccctgcgagagttcccagaagaagatcacagtcttcacctggaagttctctgaccagaggaagattctggggcagcatggcaaaggtgtattaattagaggaggttcgccttcgcagtttgatcgttttgattccaaaaaaggggcatgggagaaaggatcgtttcctctcatcatcaataaacttaagatggaagactctcagacttatatctgtgagctggagaacaggaaagaggaggtggagttgtgggtgttcaaagtgaccttcagtccgggtaccagcctgttgcaagggcagagcctgaccctgaccttggatagcaactctaaggtctctaaccccttgacagagtgcaaacacaaaaagggtaaagttgtcagtggttccaaagttctctccatgtccaacctaagggttcaggacagcgacttctggaactgcaccgtgaccctggaccagaaaaagaactggttcggcatgacactctcagtgctgggttttcagagcacagctatcacggcctataagagtgagggagagtcagcggagttctccttcccactcaactttgcagaggaaaacgggtggggagagctgatgtggaaggcagagaaggattctttcttccagccctggatctccttctccataaagaacaa agaggtgtccgtacaaaagtccaccaaagacctcaagctccagctgaaggaaacgctcccactcaccctcaagataccccaggtctcgcttcagtttgctggttctggcaacctgactctgactctggacaaagggacactgcatcaggaagtgaacctggtggtgatgaaagtggctcagctcaacaatactttgacctgtgaggtgatgggacctacctctcccaagatgagactgaccctgaagcaggagaaccaggaggccagggtctctgaggagcagaaagtagttcaagtggtggcccctgagacagggctgtggcagtgtctactgagtgaaggtgataaggtcaagatggactccaggatccaggttttatccagaggggtgaaccagacagtgttcctggcttgcgtgctgggtggctccttcggctttctgggtttccttgggctctgcatcctctgctgtgtcaggtgccggc accaacagcgccaggcagcacgaatgtctcagatcaagaggctcctcagtgagaagaagacctgccagtgcccc**<u>cac GCTatgcagGCA</u>**agccataatctcatctgataaagatcttgagtcgacaagcttgctag

#### CD4 H

acggaattccgctcgagcgccaccatggtgcgagccatctctcttaggcgcttgctgctgctgctgctgcagctgtcacaactcctagctgtcactcaagggaagacgctggtgctggggaaggaaggggaatcagcagaactgccctgcgagagttcccagaagaagatcacagtcttcacctggaagttctctgaccagaggaagattctggggcagcatggcaaaggtgtattaattagaggaggttcgccttcgcagtttgatcgttttgattccaaaaaaggggcatgggagaaaggatcgtttcctctcatcatcaataaacttaagatggaagactctcagacttatatctgtgagctggagaacaggaaagaggaggtggagttgtgggtgttcaaagtgaccttcagtccgggtaccagcctgttgcaagggcagagcctgaccctgaccttggatagcaactctaaggtctctaaccccttgacagagtgcaaacacaaaaagggtaaagttgtcagtggttccaaagttctctccatgtccaacctaagggttcaggacagcgacttctggaactgcaccgtgaccctggaccagaaaaagaactggttcggcatgacactctcagtgctgggttttcagagcacagctatcacggcctataagagtgagggagagtcagcggagttctccttcccactcaactttgcagaggaaaacgggtggggagagctgatgtggaaggcagagaaggattctttcttccagccctggatctccttctccataaagaacaaagaggtgtccgtacaaaagtccaccaaagacctcaagctccagctgaaggaaacgctcccactcaccctcaagataccccaggtctcgcttcagtttgctggttctggcaacctgactctgactctggacaaagggacactgcatcaggaagtgaacctggtggtgatgaaagtggctcagctcaacaatactttgacctgtgaggtgatgggacctacctctcccaagatgagactgaccctgaagcaggagaaccaggaggccagggtctctgaggagcagaaagtagttcaagtggtggcccctgagacagggctgtggcagtgtctactgagtgaaggtgataaggtcaagatggactccaggatccaggttttatccagaggggtgaaccagacagtgttcctggcttgcgtgctgggtggctccttcggctttctgggtttccttgggctctgcatcctctgctgtgtcaggtgccggc accaacagcgccaggcagca**<u>AACGGCCCTGGTGGCAATCCTGGTGGCAACGCAGGCGG T</u>**acctgccagtgcccccaccggatgcagaagagccataatctcatctgataaagatcttgagtcgacaagcttgctag

#### CD4 HC

acggaattccgctcgagcgccaccatggtgcgagccatctctcttaggcgcttgctgctgctgctgctgcagctgtcacaactcctagctgtcactcaagggaagacgctggtgctggggaaggaaggggaatcagcagaactgccctgcgagagttcccagaagaagatcacagtcttcacctggaagttctctgaccagaggaagattctggggcagcatggcaaaggtgtattaattagaggaggttcgccttcgcagtttgatcgttttgattccaaaaaaggggcatgggagaaaggatcgtttcctctcatcatcaataaacttaagatggaagactctcagacttatatctgtgagctggagaacaggaaagaggaggtggagttgtgggtgttcaaagtgaccttcagtccgggtaccagcctgttgcaagggcagagcctgaccctgaccttggatagcaactctaaggtctctaaccccttgacagagtgcaaacacaaaaagggtaaagttgtcagtggttccaaagttctctccatgtccaacctaagggttcaggacagcgacttctggaactgcaccgtgaccctggaccagaaaaagaactggttcggcatgacactctcagtgctgggttttcagagcacagctatcacggcctataagagtgagggagagtcagcggagttctccttcccactcaactttgcagaggaaaacgggtggggagagctgatgtggaaggcagagaaggattctttcttccagccctggatctccttctccataaagaacaaagaggtgtccgtacaaaagtccaccaaagacctcaagctccagctgaaggaaacgctcccactcaccctcaagataccccaggtctcgcttcagtttgctggttctggcaacctgactctgactctggacaaagggacactgcatcaggaagtgaacctggtggtgatgaaagtggctcagctcaacaatactttgacctgtgaggtgatgggacctacctctcccaagatgagactgaccctgaagcaggagaaccaggaggccagggtctctgaggagcagaaagtagttcaagtggtggcccctgagacagggctgtggcagtgtctactgagtgaaggtgataaggtcaagatggactccaggatccaggttttatccagaggggtgaaccagacagtgttcctggcttgcgtgctgggtggctccttcggctttctgggtttccttgggctctgcatcctctgctgtgtcaggtgccggc accaacagcgccaggcagca**<u>AACGGCCCTGGTGGCAATCCTGGTGGCAACGCAGGCGG T</u>**acc**<u>TCCcagTCC</u>**ccccaccggatgcagaagagccataatctcatctgataaagatcttgagtcgacaagcttgctag

#### CD4 LL

acggaattccgctcgagcgccaccatggtgcgagccatctctcttaggcgcttgctgctgctgctgctgcagctgtcacaactcctagctgtcactcaagggaagacgctggtgctggggaaggaaggggaatcagcagaactgccctgcgagagttcccagaagaagatcacagtcttcacctggaagttctctgaccagaggaagattctggggcagcatggcaaaggtgtattaattagaggaggttcgccttcgcagtttgatcgttttgattccaaaaaaggggcatgggagaaaggatcgtttcctctcatcatcaataaacttaagatggaagactctcagacttatatctgtgagctggagaacaggaaagaggaggtggagttgtgggtgttcaaagtgaccttcagtccgggtaccagcctgttgcaagggcagagcctgaccctgaccttggatagcaactctaaggtctctaaccccttgacagagtgcaaacacaaaaagggtaaagttgtcagtggttccaaagttctctccatgtccaacctaagggttcaggacagcgacttctggaactgcaccgtgaccctggaccagaaaaagaactggttcggcatgacactctcagtgctgggttttcagagcacagctatcacggcctataagagtgagggagagtcagcggagttctccttcccactcaactttgcagaggaaaacgggtggggagagctgatgtggaaggcagagaaggattctttcttccagccctggatctccttctccataaagaacaaagaggtgtccgtacaaaagtccaccaaagacctcaagctccagctgaaggaaacgctcccactcaccctcaagataccccaggtctcgcttcagtttgctggttctggcaacctgactctgactctggacaaagggacactgcatcaggaagtgaacctggtggtgatgaaagtggctcagctcaacaatactttgacctgtgaggtgatgggacctacctctcccaagatgagactgaccctgaagcaggagaaccaggaggccagggtctctgaggagcagaaagtagttcaagtggtggcccctgagacagggctgtggcagtgtctactgagtgaaggtgataaggtcaagatggactccaggatccaggttttatccagaggggtgaaccagacagtgttcctggcttgcgtgctgggtggctccttcggctttctgggtttccttgggctctgcatcctctgctgtgtcaggtgccggc accaacagcgccaggcagcacgaatgtctcag**<u>atcaagaggGCTGCA</u>**agtgagaagaagacctgccagtgccc ccaccggatgcagaagagccataatctcatctgataaagatcttgagtcgacaagcttgctag

#### CD4 SS

acggaattccgctcgagcgccaccatggtgcgagccatctctcttaggcgcttgctgctgctgctgctgcagctgtcacaactcctagctgtcactcaagggaagacgctggtgctggggaaggaaggggaatcagcagaactgccctgcgagagttcccagaagaagatcacagtcttcacctggaagttctctgaccagaggaagattctggggcagcatggcaaaggtgtattaattagaggaggttcgccttcgcagtttgatcgttttgattccaaaaaaggggcatgggagaaaggatcgtttcctctcatcatcaataaacttaagatggaagactctcagacttatatctgtgagctggagaacaggaaagaggaggtggagttgtgggtgttcaaagtgaccttcagtccgggtaccagcctgttgcaagggcagagcctgaccctgaccttggatagcaactctaaggtctctaaccccttgacagagtgcaaacacaaaaagggtaaagttgtcagtggttccaaagttctctccatgtccaacctaagggttcaggacagcgacttctggaactgcaccgtgaccctggaccagaaaaagaactggttcggcatgacactctcagtgctgggttttcagagcacagctatcacggcctataagagtgagggagagtcagcggagttctccttcccactcaactttgcagaggaaaacgggtggggagagctgatgtggaaggcagagaaggattctttcttccagccctggatctccttctccataaagaacaaagaggtgtccgtacaaaagtccaccaaagacctcaagctccagctgaaggaaacgctcccactcaccctcaagataccccaggtctcgcttcagtttgctggttctggcaacctgactctgactctggacaaagggacactgcatcaggaagtgaacctggtggtgatgaaagtggctcagctcaacaatactttgacctgtgaggtgatgggacctacctctcccaagatgagactgaccctgaagcaggagaaccaggaggccagggtctctgaggagcagaaagtagttcaagtggtggcccctgagacagggctgtggcagtgtctactgagtgaaggtgataaggtcaagatggactccaggatccaggttttatccagaggggtgaaccagacagtgttcctggcttgcgtgctgggtggctccttcggctttctgggtttccttgggctctgcatcctctgctgtgtcaggtgccggc accaacagcgccaggcagcacgaatg**<u>GCTcagatcaagaggctcctcGCA</u>**gagaagaagacctgccagtgcc cccaccggatgcagaagagccataatctcatctgataaagatcttgagtcgacaagcttgctag

#### CD4LL+TP

acggaattccgctcgagcgccaccatggtgcgagccatctctcttaggcgcttgctgctgctgctgctgcagctgtcacaactcctagctgtcactcaagggaagacgctggtgctggggaaggaaggggaatcagcagaactgccctgcgagagttcccagaagaagatcacagtcttcacctggaagttctctgaccagaggaagattctggggcagcatggcaaaggtgtattaatta gaggaggttcgccttcgcagtttgatcgttttgattccaaaaaaggggcatgggagaaaggatcgtttcctctcatcatcaataaacttaagatggaagactctcagacttatatctgtgagctggagaacaggaaagaggaggtggagttgtgggtgttcaaagtgaccttcagtccgggtaccagcctgttgcaagggcagagcctgaccctgaccttggatagcaactctaaggtctctaaccccttgacagagtgcaaacacaaaaagggtaaagttgtcagtggttccaaagttctctccatgtccaacctaagggttcaggacagcgacttctggaactgcaccgtgaccctggaccagaaaaagaactggttcggcatgacactctcagtgctgggttttcagagcacagctatcacggcctataagagtgagggagagtcagcggagttctccttcccactcaactttgcagaggaaaacgggtggggagagctgatgtggaaggcagagaaggattctttcttccagccctggatctccttctccataaagaacaaagaggtgtccgtacaaaagtccaccaaagacctcaagctccagctgaaggaaacgctcccactcaccctcaagataccccaggtctcgcttcagtttgctggttctggcaacctgactctgactctggacaaagggacactgcatcaggaagtgaacctggtggtgatgaaagtggctcagctcaacaatactttgacctgtgaggtgatgggacctacctctcccaagatgagactgaccctgaagcaggagaaccaggaggccagggtctctgaggagcagaaagtagttcaagtggtggcccctgagacagggctgtggcagtgtctactgagtgaaggtgataaggtcaagatggactccaggatccaggttttatccagaggggtgaaccaga cagtgttcctggcttgcgtgctgggt**<u>GTGtccttcCTC</u>**tttctgggtttccttgggctctgcatcctctgc**<u>TCTgtcaggTC C</u>**cggcaccaacagcgccaggcagcacgaatgtctcag**<u>atcaagaggGCTGCA</u>**agtgagaagaagacctgcca gtgcccccaccggatgcagaagagccataatctcatctaatgaagatcttgagtcgacaagcttgctag

#### CD4LL+R426A

acggaattccgctcgagcgccaccatggtgcgagccatctctcttaggcgcttgctgctgctgctgctgcagctgtcacaactcctagctgtcactcaagggaagacgctggtgctggggaaggaaggggaatcagcagaactgccctgcgagagttcccagaagaagatcacagtcttcacctggaagttctctgaccagaggaagattctggggcagcatggcaaaggtgtattaattagaggaggttcgccttcgcagtttgatcgttttgattccaaaaaaggggcatgggagaaaggatcgtttcctctcatcatcaataaacttaagatggaagactctcagacttatatctgtgagctggagaacaggaaagaggaggtggagttgtgggtgttcaaagtgaccttcagtccgggtaccagcctgttgcaagggcagagcctgaccctgaccttggatagcaactctaaggtctctaaccccttgacagagtgcaaacacaaaaagggtaaagttgtcagtggttccaaagttctctccatgtccaacctaagggttcaggacagcgacttctggaactgcaccgtgaccctggaccagaaaaagaactggttcggcatgacactctcagtgctgggttttcagagcacagctatcacggcctataagagtgagggagagtcagcggagttctccttcccactcaactttgcagaggaaaacgggtggggagagctgatgtggaaggcagagaaggattctttcttccagccctggatctccttctccataaagaacaaagaggtgtccgtacaaaagtccaccaaagacctcaagctccagctgaaggaaacgctcccactcaccctcaagataccccaggtctcgcttcagtttgctggttctggcaacctgactctgactctggacaaagggacactgcatcaggaagtgaacctggtggtgatgaaagtggctcagctcaacaatactttgacctgtgaggtgatgggacctacctctcccaagatgagactgaccctgaagcaggagaaccaggaggccagggtctctgaggagcagaaagtagttcaagtggtggcccctgagacagggctgtggcagtgtctactgagtgaaggtgataaggtcaagatggactccaggatccaggttttatccagaggggtgaaccaga cagtgttcctggcttgcgtgctgggtggctccttcggctttctgggtttccttgggctctgcatcctctgctgtgtcaggtgccgg**<u>c accaacagGCT</u>**caggcagcacgaatgtctcag**<u>atcaagaggGCTGCA</u>**agtgagaagaagacctgccagtgc ccccaccggatgcagaagagccataatctcatctgataaagatcttgagtcgacaagcttgctag

#### CD4LL+C

acggaattccgctcgagcgccaccatggtgcgagccatctctcttaggcgcttgctgctgctgctgctgcagctgtcacaactcctagctgtcactcaagggaagacgctggtgctggggaaggaaggggaatcagcagaactgccctgcgagagttcccagaagaagatcacagtcttcacctggaagttctctgaccagaggaagattctggggcagcatggcaaaggtgtattaattagaggaggttcgccttcgcagtttgatcgttttgattccaaaaaaggggcatgggagaaaggatcgtttcctctcatcatcaataaacttaagatggaagactctcagacttatatctgtgagctggagaacaggaaagaggaggtggagttgtgggtgttcaaagtgaccttcagtccgggtaccagcctgttgcaagggcagagcctgaccctgaccttggatagcaactctaaggtctctaaccccttgacagagtgcaaacacaaaaagggtaaagttgtcagtggttccaaagttctctccatgtccaacctaagggttcaggacagcgacttctggaactgcaccgtgaccctggaccagaaaaagaactggttcggcatgacactctcagtgctgggttttcagagcacagctatcacggcctataagagtgagggagagtcagcggagttctccttcccactcaactttgcagaggaa aacgggtggggagagctgatgtggaaggcagagaaggattctttcttccagccctggatctccttctccataaagaacaaagaggtgtccgtacaaaagtccaccaaagacctcaagctccagctgaaggaaacgctcccactcaccctcaagataccccaggtctcgcttcagtttgctggttctggcaacctgactctgactctggacaaagggacactgcatcaggaagtgaacctggtggtgatgaaagtggctcagctcaacaatactttgacctgtgaggtgatgggacctacctctcccaagatgagactgaccctgaagcaggagaaccaggaggccagggtctctgaggagcagaaagtagttcaagtggtggcccctgagacagggctgtggcagtgtctactgagtgaaggtgataaggtcaagatggactccaggatccaggttttatccagaggggtgaaccagacagtgttcctggcttgcgtgctgggtggctccttcggctttctgggtttccttgggctctgcatcctctgctgtgtcaggtgccggc accaacagcgccaggcagcacgaatgtctcag**<u>atcaagaggGCTGCA</u>**agtgagaagaagacc**<u>TCCcagTC C</u>**ccccaccggatgcagaagagccataatctcatctgataaagatcttgagtcgacaagcttgctag

The genes of CD4LL+PLXF and CD4LL+Δbind were made by cloning the in frame extraceullar domain of CD4 that encodes either the PLXF to ELXF (Glassman et al., 2018) or Δbind extracellular domain (Parrish et al., 2015) flanked by XhoI and BamHI into the CD4LL gene. The genes were sequence verified in pUC 18, subcloned into pP2 puromycin-resistance MSCV vector (MCS-IRES-puro) via 5’XhoI and 3’BglII, midi-prepped, and sequenced. The sequences for these constructs are as follows with mutated codons shown in bold uppercase letters, while the motif under interrogation is shown in bold and underlined (any non-mutated codons within the motif are bold lowercase letters and underlined):

#### CD4LL+PLXF

acggaattccgctcgagcgccaccatggtgcgagccatctctcttaggcgcttgctgctgctgctgctgcagctgtcacaactcctagctgtcactcaagggaagacgctggtgctggggaaggaaggggaatcagcagaactgccctgcgagagttcccagaagaagatcacagtcttcacctggaagttctctgaccagaggaagattctggggcagcatggcaaaggtgtattaattagaggaggttcgccttcgcagtttgatcgttttgattccaaaaaaggggcatgggagaaaggatcgtttcctctcatcatcaataaacttaagatggaagactctcagacttatatctgtgagctggagaacaggaaagaggaggtggagttgtgggtgttcaaagtgaccttcagtccgggtaccagcctgttgcaagggcagagcctgaccctgaccttggatagcaactctaaggtctctaaccccttgacagagtgcaaacacaaaaagggtaaagttgtcagtggttccaaagttctctccatgtccaacctaagggttcaggacagcgacttctggaactgcaccgtgaccctggaccagaaaaagaactggttcggcatgacactctcagtgctgggtt ttcagagcacagctatcacggcctataagagtgagggagagtcagcggagttctccttc**<u>GAActcaacGAG</u>**gcaga ggaaaacgggtggggagagctgatgtggaaggcagagaaggattctttcttccagccctggatctccttctccataaagaacaaagaggtgtccgtacaaaagtccaccaaagacctcaagctccagctgaaggaaacgctcccactcaccctcaagataccccaggtctcgcttcagtttgctggttctggcaacctgactctgactctggacaaagggacactgcatcaggaagtgaacctggtggtgatgaaagtggctcagctcaacaatactttgacctgtgaggtgatgggacctacctctcccaagatgagactgaccctgaagcaggagaaccaggaggccagggtctctgaggagcagaaagtagttcaagtggtggcccctgagacagggctgtggcagtgtctactgagtgaaggtgataaggtcaagatggactccaggatccaggttttatccagaggggtgaaccagacagtgttcctggcttgcgtgctgggtggctccttcggctttctgggtttccttgggctctgcatcctctgctgtgtcaggtg ccggcaccaacagcgccaggcagcacgaatgtctcag**<u>atcaagaggGCTGCA</u>**agtgagaagaagacctgcca gtgcccccaccggatgcagaagagccataatctcatctgataaagatcttgagtcgacaagcttgctag

#### CD4LL+Δbind

acggaattccgctcgagcgccaccatggtgcgagccatctctcttaggcgcttgctgctgctgctgctgcagctgtcacaactcctagctgtcactcaagggaagacgctggtgctggggaaggaaggggaatcagcagaactgccctgcgagagttccc agaagaagatcacagtcttcacctggaagttctctgaccagaggaagattctggggcagcatggc**<u>GATggtGATTC AGATAGC</u>**ggaggttcgccttcgcagtttgatcgttttgattccaaaaaaggggcatgggagaaaggatcgtttcctctc atcatcaataaacttaagatggaagactctcagacttatatctgtgagctggagaacaggaaagaggaggtggagttgtgggtgttcaaagtgaccttcagtccgggtaccagcctgttgcaagggcagagcctgaccctgaccttggatagcaactctaaggtctctaaccccttgacagagtgcaaacacaaaaagggtaaagttgtcagtggttccaaagttctctccatgtccaacctaagggttcaggacagcgacttctggaactgcaccgtgaccctggaccagaaaaagaactggttcggcatgacactctcagtgctgggttttcagagcacagctatcacggcctataagagtgagggagagtcagcggagttctccttcccactcaactttgcagaggaaaacgggtggggagagctgatgtggaaggcagagaaggattctttcttccagccctggatctccttctccataaagaacaaagaggtgtccgtacaaaagtccaccaaagacctcaagctccagctgaaggaaacgctcccactcaccctcaagataccccaggtctcgcttcagtttgctggttctggcaacctgactctgactctggacaaagggacactgcatcaggaagtgaacctggtggtgatgaaagtggctcagctcaacaatactttgacctgtgaggtgatgggacctacctctcccaagatgagactgaccctgaagcaggagaaccaggaggccagggtctctgaggagcagaaagtagttcaagtggtggcccctgagacagggctgtggcagtgtctactgagtgaaggtgataaggtcaagatggactccaggatccaggttttatccagaggggtgaaccagacagtgttcctggcttgcgtgctgggtggctccttcggctttctgggtttccttgggctctgcatcctctgctgtgtca ggtgccggcaccaacagcgccaggcagcacgaatgtctcag**<u>atcaagaggGCTGCA</u>**agtgagaagaagacctg ccagtgcccccaccggatgcagaagagccataatctcatctgataaagatcttgagtcgacaagcttgctag

## Acknowledgements

This work was supported by the National Institutes of Health/National Institute of Allergy and Infections Diseases Grant R01AI101053 (MSK), the Pew Scholars Program in the Biomedical Sciences (MSK), the Cancer Center Support Grant CCSG-CA 023074 for flow cytometry (MSK), and AZ TRIF funds from the BIO5 institute (KVD). We thank Thomas Serwold for critical feedback as well as Dominik Schenten, Joseph Harrison, Piet Maes for critically reading the manuscript.

## Author’s Contributions

MSK and KVD conceived of the project. MSK directed the research and KVD performed the computational analysis. MSL, PJT, CK, KL and HLP performed the experiments. The manuscript was written by MSK and KVD. All authors contributed to data analysis, discussions, editing, reading and approval of the manuscript.

---

**Figure S1.**
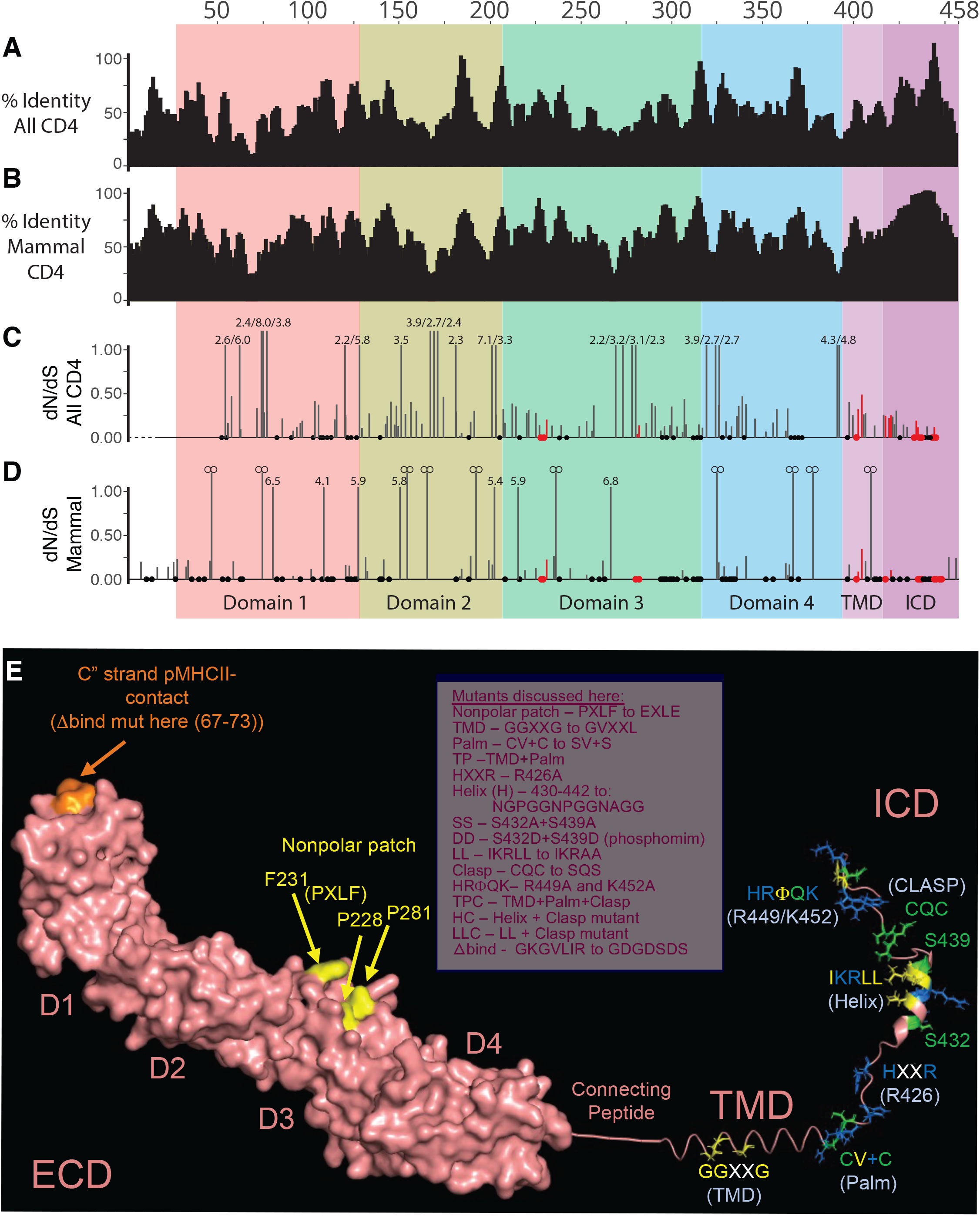
Conservation and purifying selection of the CD4 molecule. (A) Mean pairwise sequence identity over all pairs in the alignment column was calculated using a sliding window of size 5. The alignment was based on 99 CD4 orthologs and still contains the sequences downstream of the CQC clasp. These residues were later removed based on low sequence conservation. (B) As in (A) but including only mammalian CD4 orthologs. (C) The Fixed Effects Likelihood (FEL) model was used to estimate non-synonymous (dN) and synonymous (dS) substitution rates on a per-site basis as an indication of evolutionary selection. Bars show the dN/dS ratio of both these values. Only bars for which the likelihood ratio test indicated statistical significance (P < 0.1) are shown. dN/dS ratios below 1 are indicative of purifying selection, while a ratio great than 1 are under positive selection. For visualization purposes, we focus on values lower than 1. For codons under positive selection (i.e, > 1), the value indicates the dN/dS ratio. An infinity symbol was used to indicate those codons for which dS = 0. Circles have a dN/dS ration of 0. Red highlighted codons are located within motifs identified within this study. CD4 domains are indicated using color. Numbering according to mouse CD4. (E) A theoretical structural model to show the relative location of the motifs discussed here. The surface rendered ECD (pdb 1WIQ), joined with a connecting peptide and TMD (built using the PyMol Molecular Graphics system), and ICD (pdb 1Q68). See also Figure 1.

**Figure S2.**
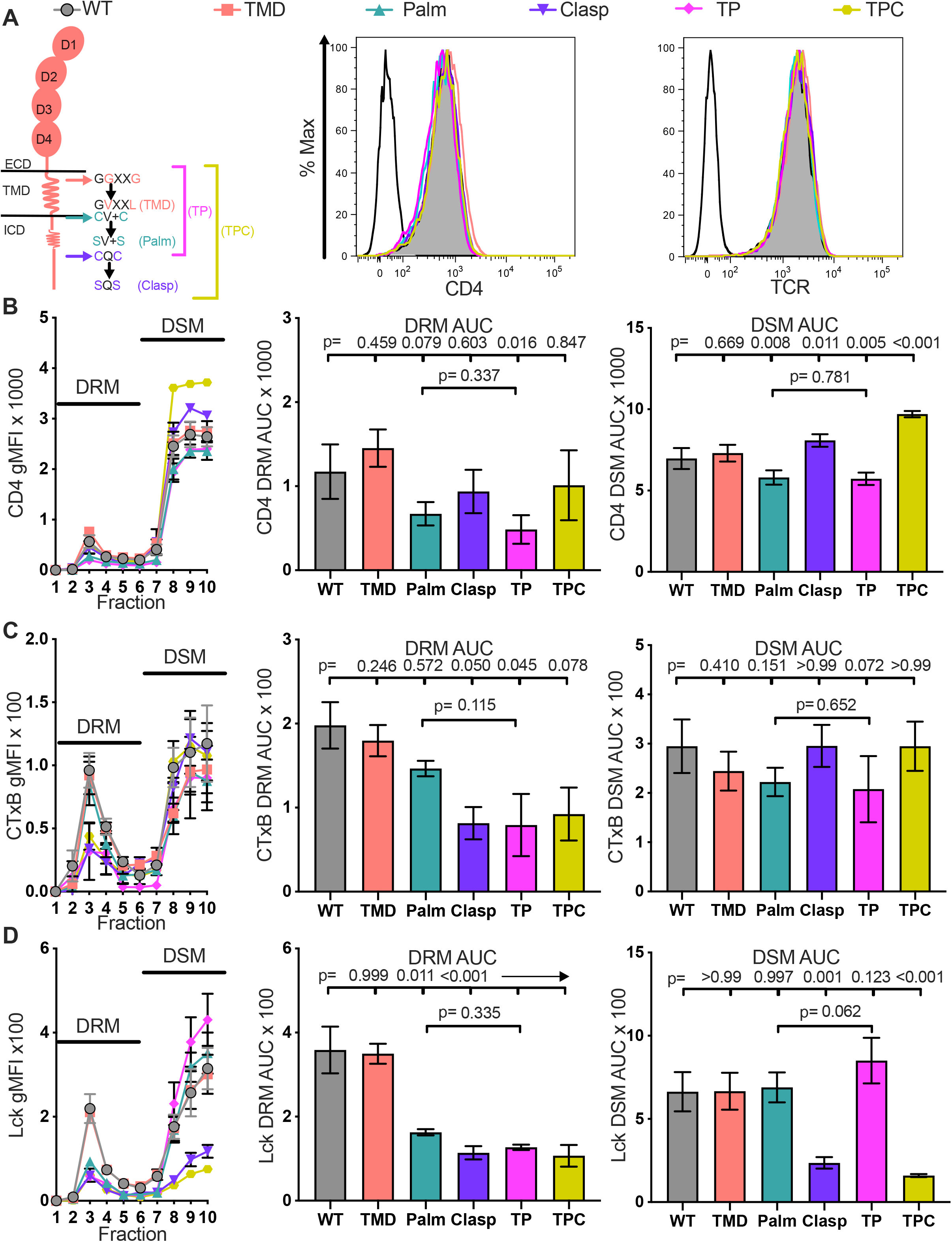
Analysis of TMD, Palm, Clasp, TP, and TPC mutant cell lines. (A) Cartoon of CD4 highlighting the location and nature of the mutations tested (left). Flow cytometry analysis of CD4 (center) and TCR expression (right) on 58α^-^β^-^ cells. (B) Raw CD4 signal data is shown for each sucrose fraction (left). AUC is shown for the DRM fractions (center) and DSM fractions (right). (C) Raw cholera toxin B (CTxB) signal data is shown for each sucrose fraction (left). AUC is shown for the DRM (center) and DSM fractions (right). (D) Raw Lck signal data is shown for each sucrose fraction (left). AUC is shown for the DRM (center) and DSM fractions (right). For (B-D) each data point represents the mean +/- SEM for the same three independent experiments (biological replicates). One-way ANOVA was performed with a Dunnett’s posttest for comparisons with WT samples, and a Sidak’s posttest for comparisons between selected samples. See also Figure 2.

**Figure S3.**
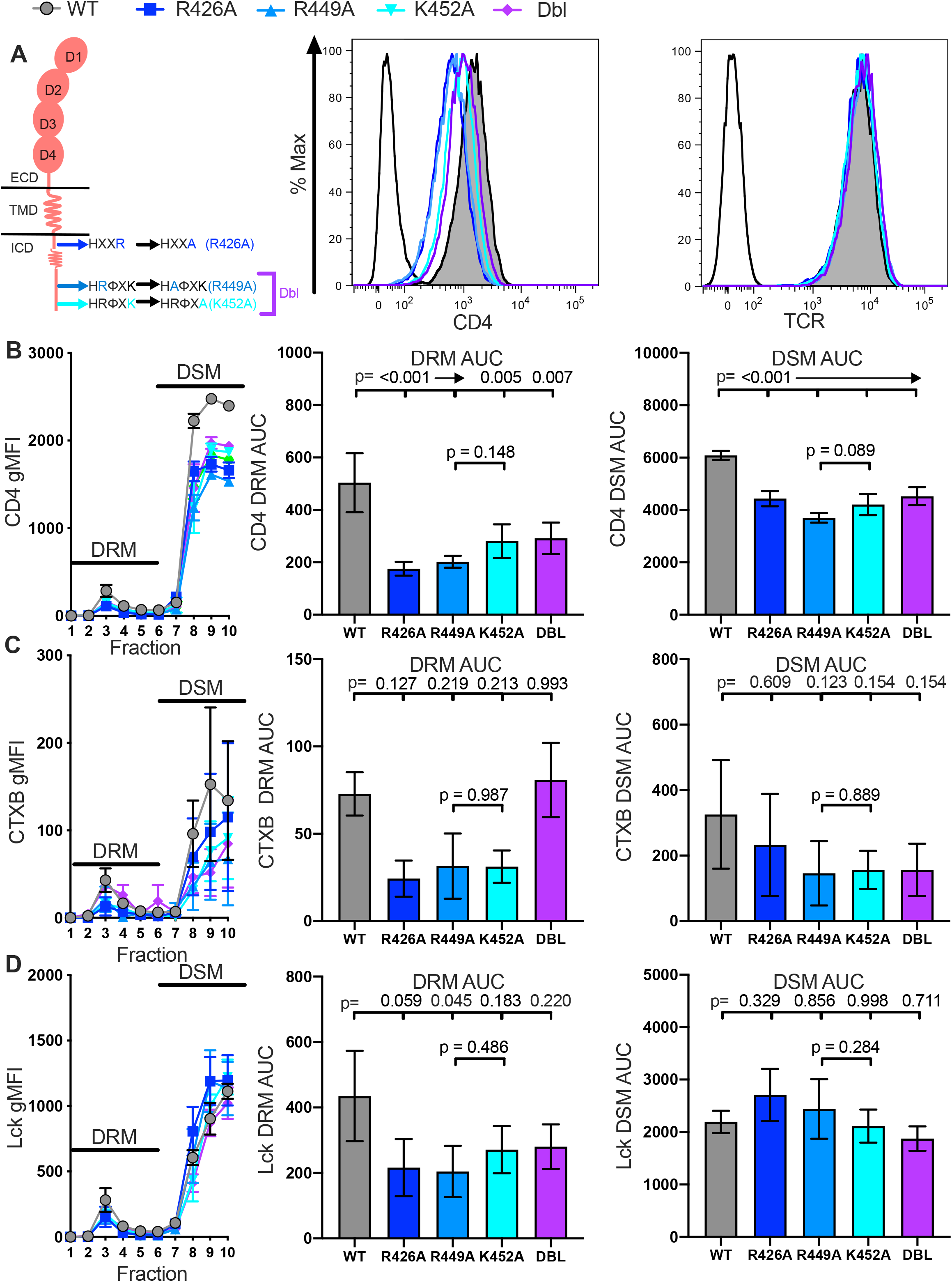
Analysis of R426A, R449A, K452A, and Dbl mutant cell lines. (A) Cartoon of CD4 highlighting the location and nature of the mutations tested (left). Flow cytometry analysis of CD4 (center) and TCR expression (right) on 58α^-^β^-^ cells. (B) Raw CD4 signal data is shown for each sucrose fraction (left). AUC is shown for the DRM fractions (center) and DSM fractions (right). (C) Raw cholera toxin B (CTxB) signal data is shown for each sucrose fraction (left). AUC is shown for the DRM (center) and DSM fractions (right). (D) Raw Lck signal data is shown for each sucrose fraction (left). AUC is shown for the DRM (center) and DSM fractions (right). For (B-D) each data point represents the mean +/- SEM for the same three independent experiments (biological replicates). One-way ANOVA was performed with a Dunnett’s posttest for comparisons with WT samples, and a Sidak’s posttest for comparisons between selected samples. See also Figure 3.

**Figure S4.**
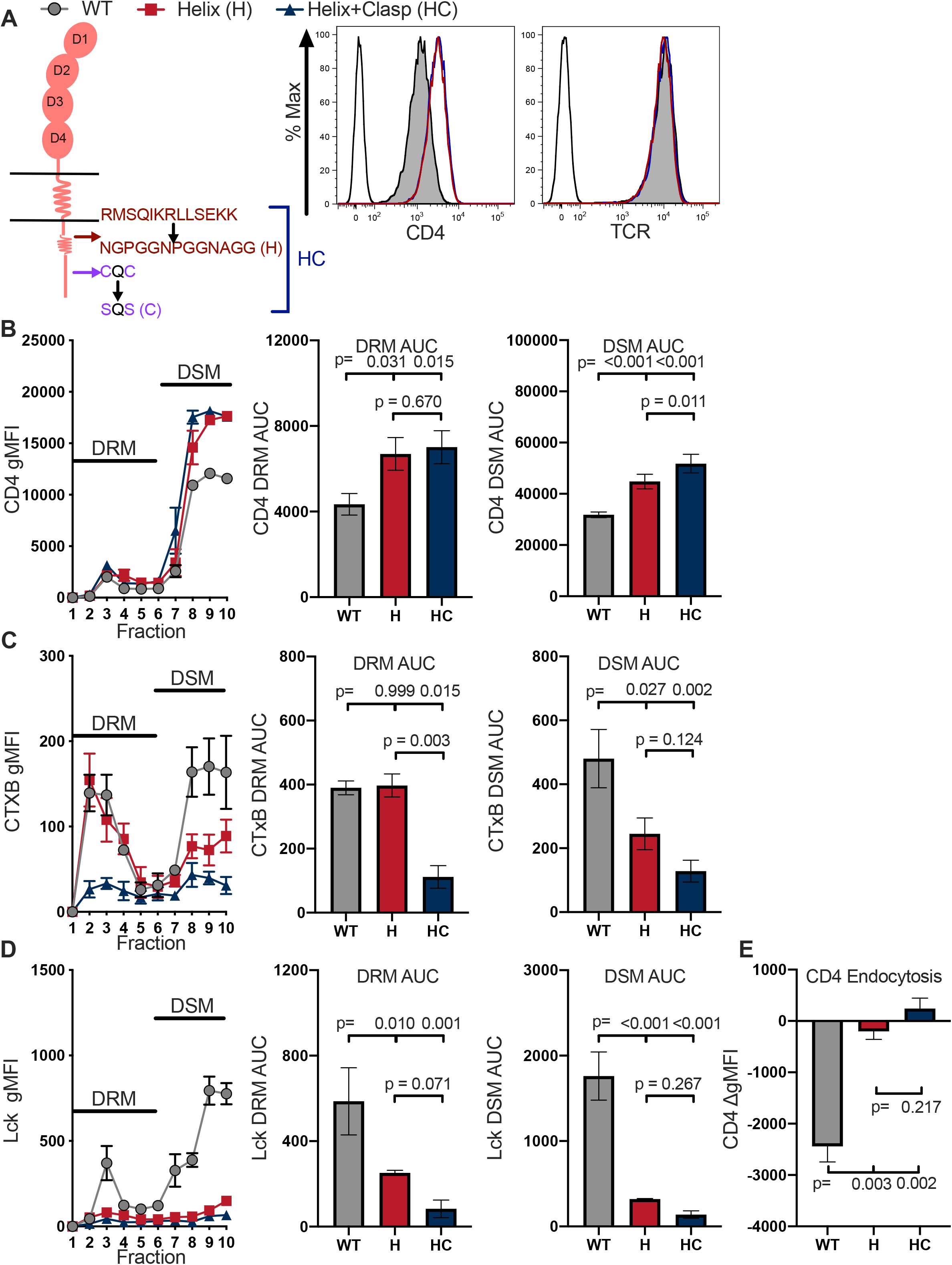
Analysis of H and HC mutant cell lines. (A) Cartoon of CD4 highlighting the location and nature of the mutations tested (left). Flow cytometry analysis of CD4 (center) and TCR expression (right) on 58α^-^β^-^ cells. (B) Raw CD4 signal data is shown for each sucrose fraction (left). AUC is shown for the DRM fractions (center) and DSM fractions (right). (C) Raw cholera toxin B (CTxB) signal data is shown for each sucrose fraction (left). AUC is shown for the DRM (center) and DSM fractions (right). (D) Raw Lck signal data is shown for each sucrose fraction (left). AUC is shown for the DRM (center) and DSM fractions (right). (E) CD4 endocytosis after pMHCII engagement is shown for the indicated cell lines after 16 hours coculture with APCs in the presence of 10µM MCC peptide. The change in CD4 gMFI, as measured by flow cytometry, is shown for each cell line relative to an equivalent sample cultured with APCs in the absence of MCC peptide. For (B-E) each data point represents the mean +/- SEM for the same three independent experiments (biological replicates). For E, endocytosis measurements were performed in triplicate (technical replicates) for each experiment. One-way ANOVA was performed with a Dunnett’s posttest for comparisons with WT samples, and a Sidak’s posttest for comparisons between selected samples. See also Figure 4.

**Figure S5.**
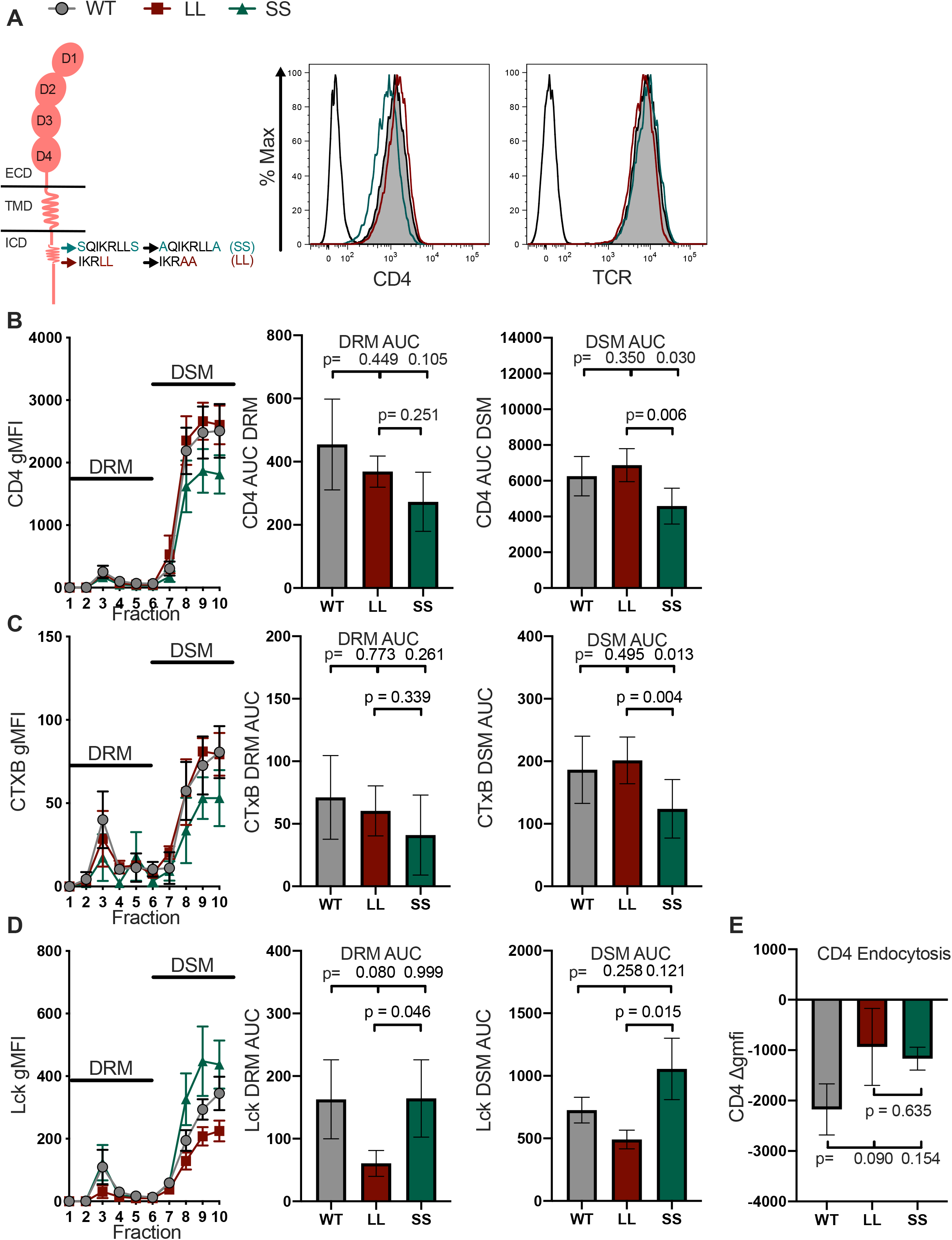
Analysis of LL and SS cell lines. (A) Cartoon of CD4 highlighting the location and nature of the mutations tested (left). Flow cytometry analysis of CD4 (center) and TCR expression (right) on 58α^-^β^-^ cells. (B) Raw CD4 signal is shown for each sucrose fraction (left). AUC is shown for the DRM fractions (center) and DSM fractions (right). (C) Raw cholera toxin B (CTxB) signal is shown for each sucrose fraction (left). AUC is shown for the DRM (center) and DSM fractions (right). (D) Raw Lck signal is shown for each sucrose fraction (left). AUC is shown for the DRM (center) and DSM fractions (right). (E) CD4 endocytosis after pMHCII engagement is shown for the indicated cell lines after 16 hours coculture with APCs in the presence of 10µM MCC peptide. The change in CD4 gMFI, as measured by flow cytometry, is shown for each cell line relative to an equivalent sample cultured with APCs in the absence of MCC peptide. For (B-E) each data point represents the mean +/- SEM for the same three independent experiments (biological replicates). For E, endocytosis measurements were performed in triplicate (technical replicates) for each experiment. One-way ANOVA was performed with a Dunnett’s posttest for comparisons with WT samples, and a Sidak’s posttest for comparisons between selected samples. See also Figure 5.

**Figure S6.**
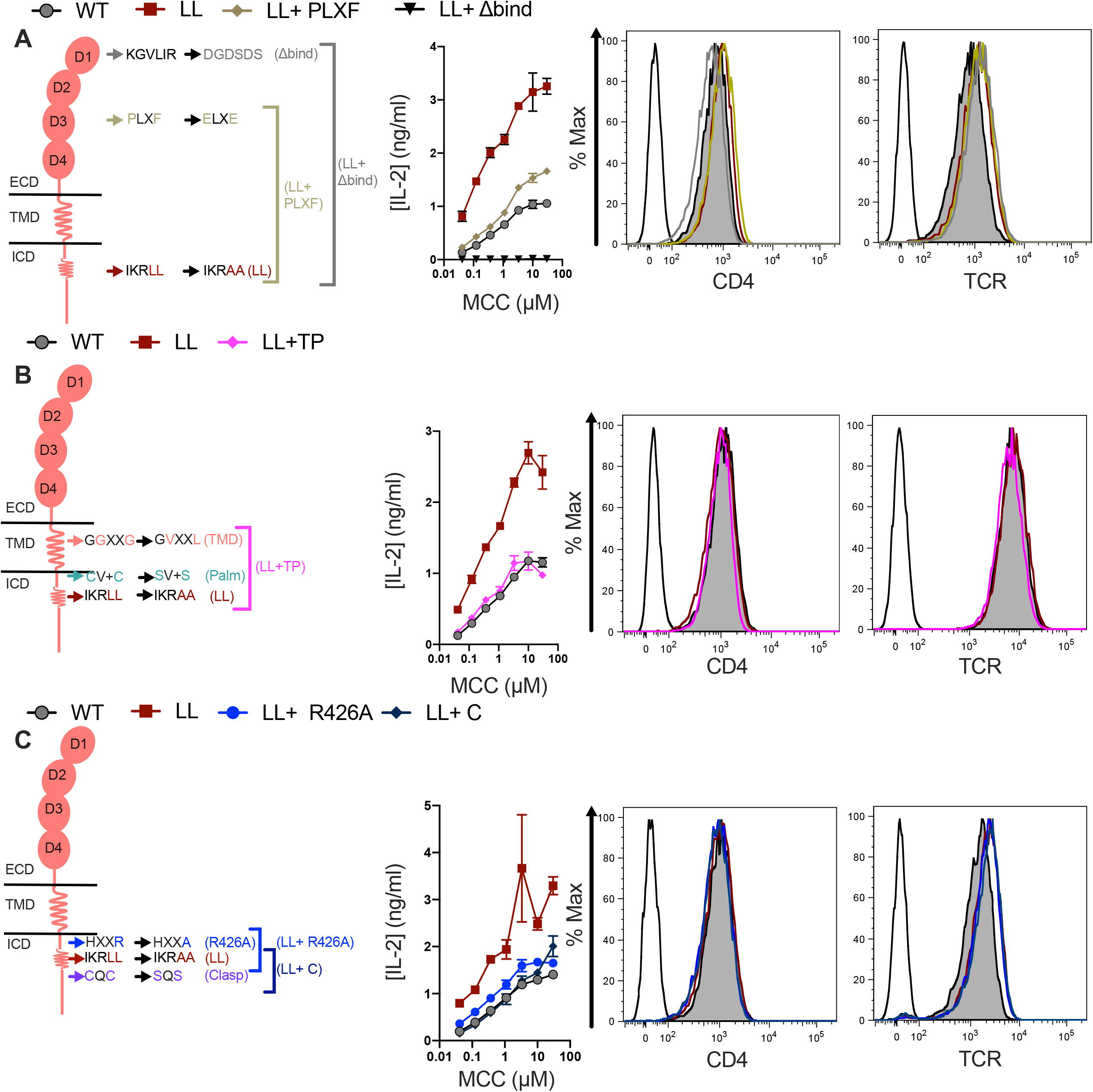
Analysis of CD4 mutants of coevolving motifs. (A-D) Cartoon of CD4 highlighting the location and nature of the mutations tested (left). Representative IL-2 dose responses to MCC peptide for one of three independent experiments with each dataset representing the mean +/- SEM of triplicates (center). Flow cytometry analysis of CD4 and TCR expression on 58α^-^β^-^ cells (right). See also Figure 7.

